# Genetic predisposition to mosaic Y chromosome loss in blood is associated with genomic instability in other tissues and susceptibility to non-haematological cancers

**DOI:** 10.1101/514026

**Authors:** Deborah J. Thompson, Giulio Genovese, Jonatan Halvardson, Jacob C. Ulirsch, Daniel J. Wright, Chikashi Terao, Olafur B. Davidsson, Felix R. Day, Patrick Sulem, Yunxuan Jiang, Marcus Danielsson, Hanna Davies, Joe Dennis, Malcolm G. Dunlop, Douglas F. Easton, Victoria A. Fisher, Florian Zink, Richard S. Houlston, Martin Ingelsson, Siddhartha Kar, Nicola D. Kerrison, Ragnar P. Kristjansson, Rong Li, Chey Loveday, Jonas Mattisson, Steven A. McCarroll, Yoshinori Murakami, Anna Murray, Pawel Olszewski, Edyta Rychlicka-Buniowska, Robert A. Scott, Unnur Thorsteinsdottir, Ian Tomlinson, Behrooz Torabi Moghadam, Clare Turnbull, Nicholas J. Wareham, Daniel F. Gudbjartsson, INTEGRAL-ILCCO, The Breast Cancer Association Consortium, CIMBA, The Endometrial Cancer Association Consortium, The Ovarian Cancer Association Consortium, The PRACTICAL Consortium, The Kidney Cancer GWAS Meta-Analysis Project, eQTLGen Consortium, BIOS Consortium, 23andMe Research Team, Yoichiro Kamatani, Hilary K. Finucane, Eva R. Hoffmann, Steve P. Jackson, Kari Stefansson, Adam Auton, Ken K. Ong, Mitchell J. Machiela, Po-Ru Loh, Jan P. Dumanski, Stephen J. Chanock, Lars A. Forsberg, John R. B. Perry

## Abstract

Mosaic loss of chromosome Y (LOY) in circulating white blood cells is the most common form of clonal mosaicism, yet our knowledge of the causes and consequences of this is limited. Using a newly developed approach, we estimate that 20% of the UK Biobank male population (N=205,011) has detectable LOY. We identify 156 autosomal genetic determinants of LOY, which we replicate in 757,114 men of European and Japanese ancestry. These loci highlight genes involved in cell-cycle regulation, cancer susceptibility, somatic drivers of tumour growth and cancer therapy targets. Genetic susceptibility to LOY is associated with non-haematological health outcomes in both men and women, supporting the hypothesis that clonal haematopoiesis is a biomarker of genome instability in other tissues. Single-cell RNA sequencing identifies dysregulated autosomal gene expression in leukocytes with LOY, providing insights into how LOY may confer cellular growth advantage. Collectively, these data highlight the utility of studying clonal mosaicism to uncover fundamental mechanisms underlying cancer and other ageing-related diseases.

## Introduction

Each day the human body produces billions of highly specialised blood cells, generated from a self-renewing pool of 50,000-200,000 haematopoietic stem cells (HSCs)^1^. As these cells age and divide, mutation and mitotic errors create genetic diversity within the HSC pool and their progenitors. If a genetic alteration confers a selective growth advantage to one cell over the others, clonal expansion may occur. This process propels the lineage to disproportionate frequency, creating a genetically distinct sub-population of cells. In the literature this is commonly referred to as clonal haematopoiesis, or more broadly (not restricting to considering leukocytes), clonal mosaicism^2^ or aberrant clonal expansion^3^.

Population-based studies assessing the magnitude and effect of clonal mosaicism have been largely limited by the challenges of accurately detecting the expected low cell-fraction mosaic events in leukocytes using genotype-array or sequence read data^4^. Recent advances in statistical methodology have improved sensitivity, with approaches now able to catalogue mosaic events at higher resolution across the genome^5,6^. Detection of large structural mosaic events can vary considerably in size – from 50kb to entire chromosomes in length – and are typically present in only a small fraction of circulating leukocytes (<5%). It is well established that loss of the sex chromosomes – particularly the Y chromosome (LOY) in men – is by far the most frequently observed somatic change in leukocytes^7–9^. It remains unclear if and why absence of a Y chromosome provides a selective growth advantage in these cells – we hypothesise this could be due to (amongst other unknown mechanisms) the loss of a putative Y-linked cell-growth suppressor gene, loss of a Y-linked transcription factor influencing expression of cell-growth related autosomal genes or the reduced energy cost of cellular divisions.

Our understanding of why some individuals, but not others, exhibit clonal mosaicism in blood is also limited. Previous studies have demonstrated robust associations with age, sex (clonal mosaicism is more frequent in males), smoking and inherited germline genetic predisposition^2,4,10–15^. Recent epidemiological studies have challenged the view that LOY in the hematopoietic system is a phenotypically neutral event, with epidemiological associations observed with various forms of cancer^13,16–20^, autoimmune conditions^21,22^, age-related macular degeneration^23^, cardiovascular disease^24^, Alzheimer’s disease^25^, type 2 diabetes^15^, obesity^15^, and all-cause mortality^15,16^. The extent to which such observations represent a causal association, reverse causality or confounding is unclear. Furthermore, if these do represent causal effects, the mechanisms underlying such effects are unknown.

Key questions are whether loss of a Y chromosome from circulating leukocytes has a direct functional effect (for example, impairs immune function) and whether LOY in leukocytes is a barometer of broader genomic instability in other cell types. Understanding the mechanisms that drive clonal mosaicism and identifying genes which promote proliferative advantage to cells may help answer these questions and provide important insights into mechanisms of diseases of ageing. To this end we sought to identify novel susceptibility loci for LOY, an attractive form of clonal mosaicism to study given its relative ease of detection and high prevalence in the male population. Previous genome-wide association studies (GWAS) for LOY identified 19 common susceptibility loci and highlighted its relevance as a biomarker of cell cycle efficiency and DNA damage response (DDR) in leukocytes^13,14^. Here, we develop and apply a new computational approach for detecting LOY to over 200,000 men from the UK Biobank study. We identify 137 novel loci which we use, along with the known 19 loci^14^, to demonstrate a shared genetic architecture between LOY, non-haematological cancer susceptibility and reproductive ageing in women. These data, in aggregate, support the hypothesis that LOY in leukocytes is a biomarker of genomic instability in other cell types with functional consequences across diverse biological systems.

## Results

Previous studies assessing LOY have used a quantitative measure derived from the average intensity log-R ratio (termed mLRR-Y) of all array-genotyped Y chromosome single-nucleotide polymorphisms (SNPs). Here, we adapted a recently developed long-range phasing approach for mosaic event detection to estimate a dichotomous classification, which uses allele-specific genotyping intensities in the pseudo-autosomal region (we term this PAR-LOY, see Methods). This was applied to 205,011 men from UKBB (aged 40-70) in whom we identified 41,791 (20%) with detectable LOY. Men classified as LOY had an mLRR-Y score (derived using variants outside of the PAR) 0.9 standard deviations lower on average (95% CI 0-88-0.9) than non-LOY males (mean mLRR-Y −0.046 vs 0.009), reflecting the expected lower level of intensity due to reduced Y chromosome genetic material. Consistent with previous observations of clonal mosaicism, current smokers were at a higher risk of LOY (odds ratio (OR) 1.62 [95% CI 1.57-1.66]) and there was a strong association with age; the prevalence increased from 2.5% at age 40 to 43.6% at age 70 (Figure 1).

**Figure 1.**
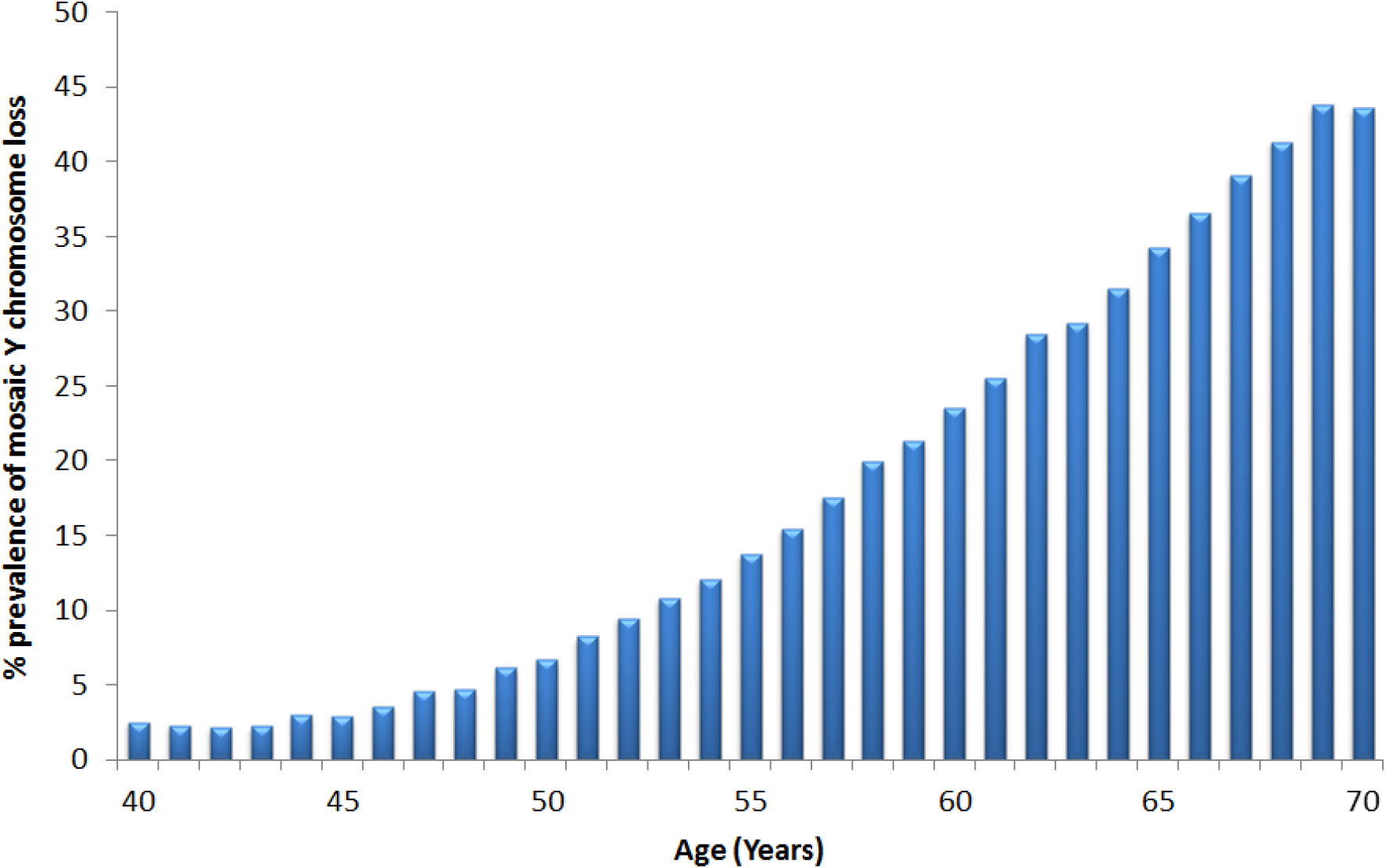
Prevalence of mosaic Y chromosome loss by age in UK Biobank study participants

### The genetic architecture of mosaic Y chromosome loss

We estimated a heritability of 31.7% (95CI 29.9 to 33.6%) for LOY, distributed across all individual chromosomes in proportion to their relative sizes (**Figure S1**). To identify individual genetic variants underlying this heritability we performed a GWAS for LOY, identifying 18,146 variants with genome-wide significant associations (P<5×10^−8^). We resolved these into 156 statistically independent signals (**Table S1**), which included all 19 loci previously reported^14^. Effect sizes for these 156 associations ranged from OR 1.03-2.02, with LOY risk allele frequencies between 0.25% and 99.8% (Figure 2).

**Figure 2.**
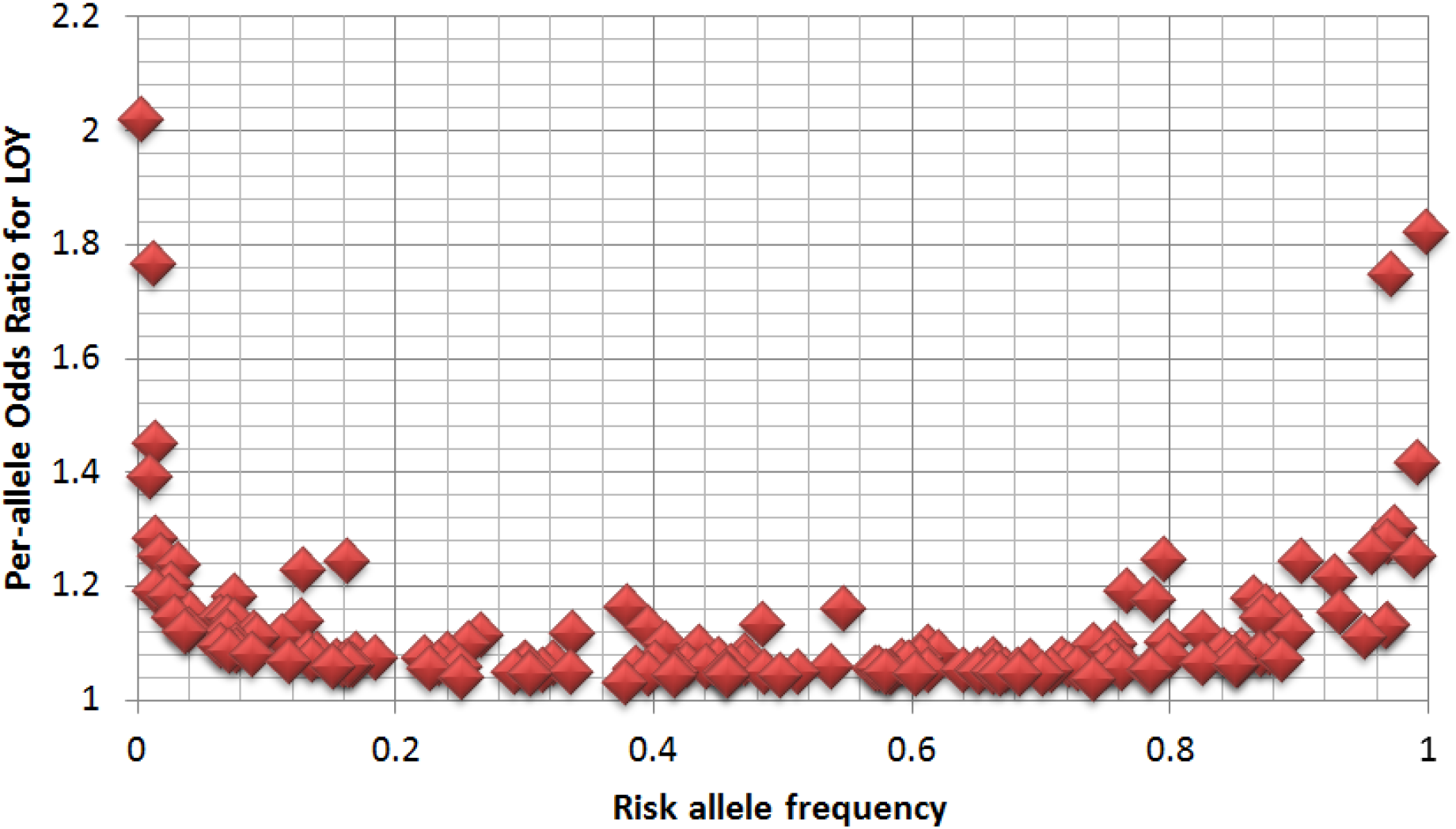
Distribution of allele frequency and effect size for the 156 identified LOY loci

We directly compared the power of our PAR-LOY calls to the previously used mLRR-Y derived measures by performing an mLRR-Y based GWAS in the same current study samples (**Table S1**). Across the 156 loci we observed an average ~2.5x increase in χ^2^ association statistic, exemplified by the strongest associated variant (rs17758695-*BCL2*) increasing in significance from P_mLRRy_=7.5×10^−65^ to P_PAR-LOY=_4.1×10^−147^. Only 61 of the 156 loci would have reached genome-wide significance in an mLRR-Y based analysis. Across the genome the lambda GC (ratio of expected to observed median test statistic) increased from 1.15 to 1.20 (mean χ^2^ from 1.28 to 1.47), with no evidence of signal inflation due to population structure (LD score regression intercept 1.01).

To confirm the validity of our identified signals we sought replication in three independent datasets. Firstly, we used data generated using 653,019 male research participants from the personal genetics company 23andMe, Inc. (**Table S1**). These samples differed from the discovery samples both in terms of DNA source (saliva rather than peripheral blood) and LOY measurement type (quantitative mLRR-Y rather than dichotomous PAR-LOY calls). Despite this heterogeneity, all but one of the 154 loci (2 failed QC) had directionally concordant effects (binomial sign test P=1.4×10^−44^), with 126 exhibiting nominally significant association (P<0.05) and 88 at a more conservative threshold (P<0.05/156). Secondly, we sought further confirmation from the Icelandic deCODE study (N=8,715) where LOY was estimated using sequence reads from whole genome sequencing (DNA extracted from blood), rather than array data. These data demonstrated an overall directional consistency of 94% across the associated loci (140/149 variants tested, binomial sign test P=2.3×10^−31^) and 74 nominal associations (**Table S1**). Third, we replicated our loci in a set of 95,380 Japanese ancestry men from the BBJ project, with LOY estimated using mLRR-Y in whole blood. Of the 100/156 variants which passed QC and were polymorphic in East Asians, 92 had a consistent direction of effect (binomial P=3.2×10^−19^). Of these, 29 reached genome-wide significance in these data alone and 73 had at least nominal association (**Table S1**).

Finally, a negative control analysis using mLRR-Y estimated in 245,349 UKBB women (**Table S1**) – reflecting experimental noise in intensity variation – did not produce any significant associations after Bonferroni correction across the 156 loci (P_max_=4.3×10^−3^). In aggregate, these data strongly suggest that our discovery analysis identifies genetic determinants of LOY that are robust to ancestry, measurement technique and DNA source.

### Implicated genes, cell types and biological pathways

We used various approaches to move from genomic association to identifying potentially causal variants, functional genes, cell types and biological pathways associated with LOY (see Methods). First, we performed Bayesian fine-mapping (see Methods) to quantify the probability that any single variant at a locus was causal for LOY by disentangling the effects of linkage disequilibrium (LD) (Figure 3, **Table S2-S3**). Fine-mapping identified at least one variant with reasonable confidence (posterior probability [PP] > 10%) in 80% (101/126) of regions, including at least one very high confidence variant (PP > 75%) in 25% (31/126) of regions (Figure 3A). These variants were enriched in exons of protein coding genes, their promoters, their transcribed but untranslated regions, and in hematopoietic regulatory regions marked by accessible chromatin (Figure 3B, **Table S4**).

**Figure 3.**
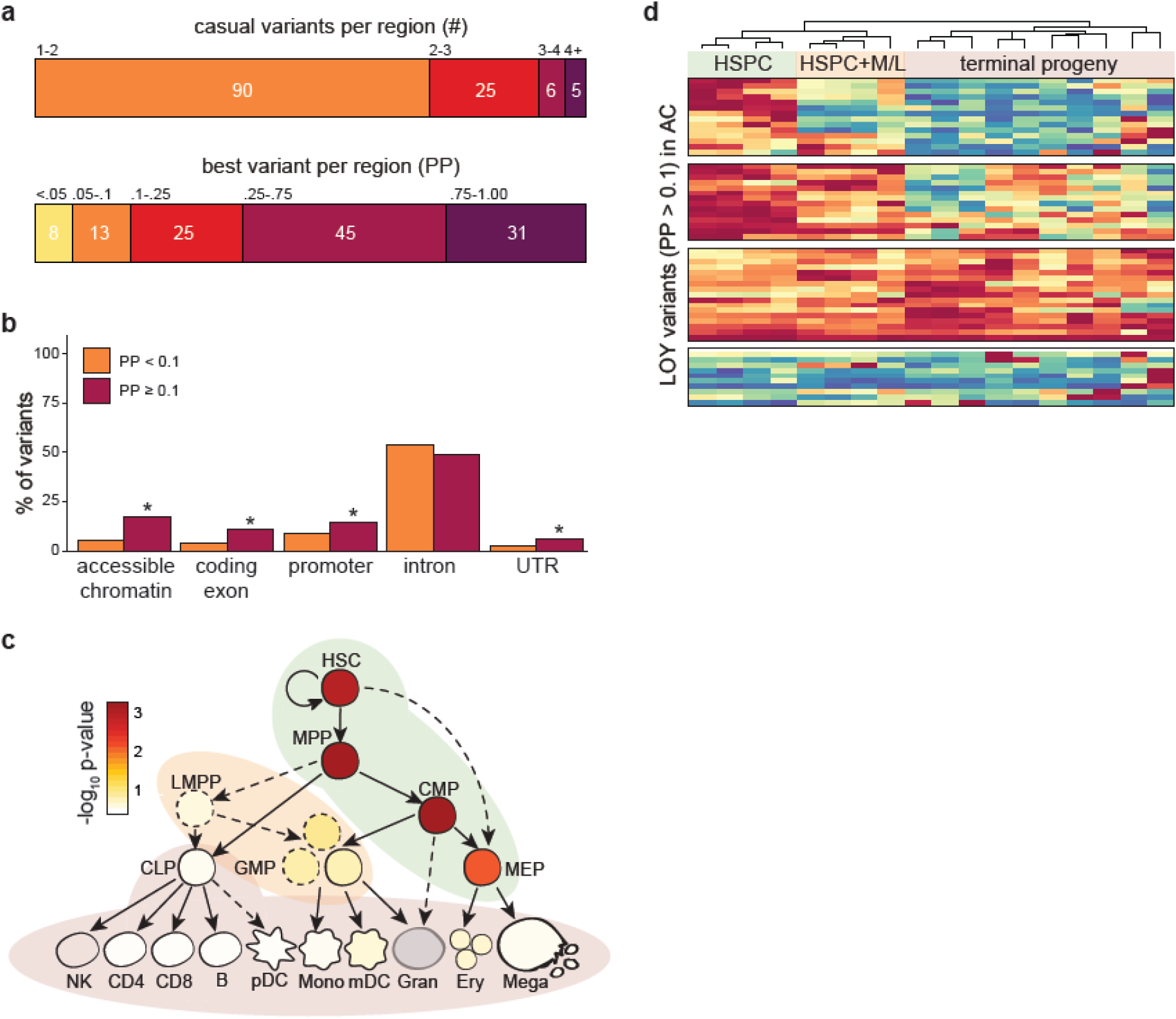
Results from fine-mapping analyses. Panel a shows the posterior expected number of causal variants (*top*) as well as the best fine-mapped variant (*bottom*) in each region. Genomic enrichments for variants stratified by posterior probability are shown in panel **b.** Fine-mapped variants were enriched for accessible chromatin in hematopoiesis, as well as in exons, promoters, and UTRs of protein coding genes, but not for introns. Panel c shows g-chromVAR cell-type enrichments across the hematopoietic tree for LOY. HSCs, MPPs, and CMPs meet Bonferroni threshold (α = 0.05 / 18). Developmental patterns of accessible chromatin for variants with posterior probability > 10% are shown in panel **d**, revealing that 14 variants are fully restricted to acting within HSPCs, 14 variants can also have regulatory effects in myeloid and lymphocyte progenitors, and 17 variants are capable of acting across the majority of hematopoiesis. K-means clustering (k = 4 determined by the gap statistic) was used to identify patterns of accessibility, and cell types were hierarchically clustered. HSC, hematopoietic stem cell; MPP, multi-potent progenitor; CMP, common myeloid progenitor; HSPC, hematopoietic stem and progenitor cell; M/L, myeloid and lymphoid; PP, posterior probability; AC, accessible chromatin; UTR, untranslated region; PChiC, promoter capture Hi-C; eQTL, expression quantitative trait locus; corr, ATAC/chromatin-RNA correlations

Using both fine-mapped variants and genome-wide polygenic signal (see Methods), we found that hematopoietic stem and progenitor cells (HSPCs) were the most strongly enriched cell-types for LOY associated variants (Figure 3C, Figure S2, Table S5, Table S6). Amongst the fine-mapped variants, we further subdivided this enrichment into 3 distinct temporal modes indicative of increasing regulatory capacity across haematopoiesis (Figure 3D). These observations suggest that many of our identified variants exert their effects directly in hematopoietic stem cells, rather than further differentiated white blood cell types. This is in stark contrast to variants associated with the production of terminal blood cell types, which are enriched at terminal blood progenitors and depleted in HSPCs^26^.

We next used two approaches (see methods) to map associated genetic variants to genes via expression effects (eQTLs) in whole blood, implicating a total of 110 unique transcripts (**Table S8-S10**). This included the *HLA-A* gene, where our lead variant in this region (6:29835518_T_A) tagged the HLA-A*02:01 allele (**Table S11**). We also identified genes harbouring a non-synonymous variant either fine-mapped (PP > 10%) or in high LD (r^2^ > 0.8) with an index variant, highlighting 22 genes (**Table S8**).

Biological pathway analysis using two approaches (see methods) identified a number of associated pathways, the majority of which converged on aspects of cell cycle regulation and DNA damage response (**Table S12 and S13**).

### Overlap between LOY associated variants and cancer susceptibility loci

While detectable clonal mosaicism is clearly associated with future risk of haematological cancers^6^, its relationship with other cancers is less clear. Using data curated by the Open Targets platform (see Methods), we found that LOY-associated variants were preferentially found near genes involved in cancer susceptibility (P=9.9×10^−7^), somatic drivers of tumour growth (P=7×10^−4^) and targets of approved or in trial cancer therapies (P=0.05). In total, 18 of the 156 mosaic leukocyte LOY associated variants were correlated (r^2^>0.1) with known susceptibility variants for one or more type of non-haematological cancer (**Table S14**), including breast, prostate, testicular, kidney, melanoma and brain. Notable examples include a loss-of-function variant in *CHEK2* (rs186430430 r^2^~1 with frameshift variant 1100delC) which confers a ~2.3 fold high risk of breast cancer^27^, and an intronic signal (rs56345976) in the telomerase reverse transcriptase (*TERT*) gene which is in modest LD (r^2^~0.12) with variants associated with longer telomeres and with increased risks of breast, ovarian, prostate cancers and glioblastoma, but also seen to be protective in other cancers^28^.

To systematically assess the relationship between LOY susceptibility and cancer risk, we tested a genetic risk score comprised of our 156 variants on two male-specific cancers (Figure 4, **Table S15**). Genetically-predicted LOY was associated with both increased risk of prostate cancer (OR=1.68 95% CI 1.33-2.11, P=1.9×10^−5^) and testicular germ cell tumour (OR 2.97 (1.45-6.07) P=0.003). Additional publicly available GWAS data for cancers in both sexes showed directionally consistent associations for renal cell carcinoma (OR 1.12 (1.04-1.21) P=0.004), lung cancer (OR 1.28 (0.98-1.68), P=0.07) and colorectal cancer (OR 1.18 (0.93-1.50), P=0.16).

**Figure 4.**
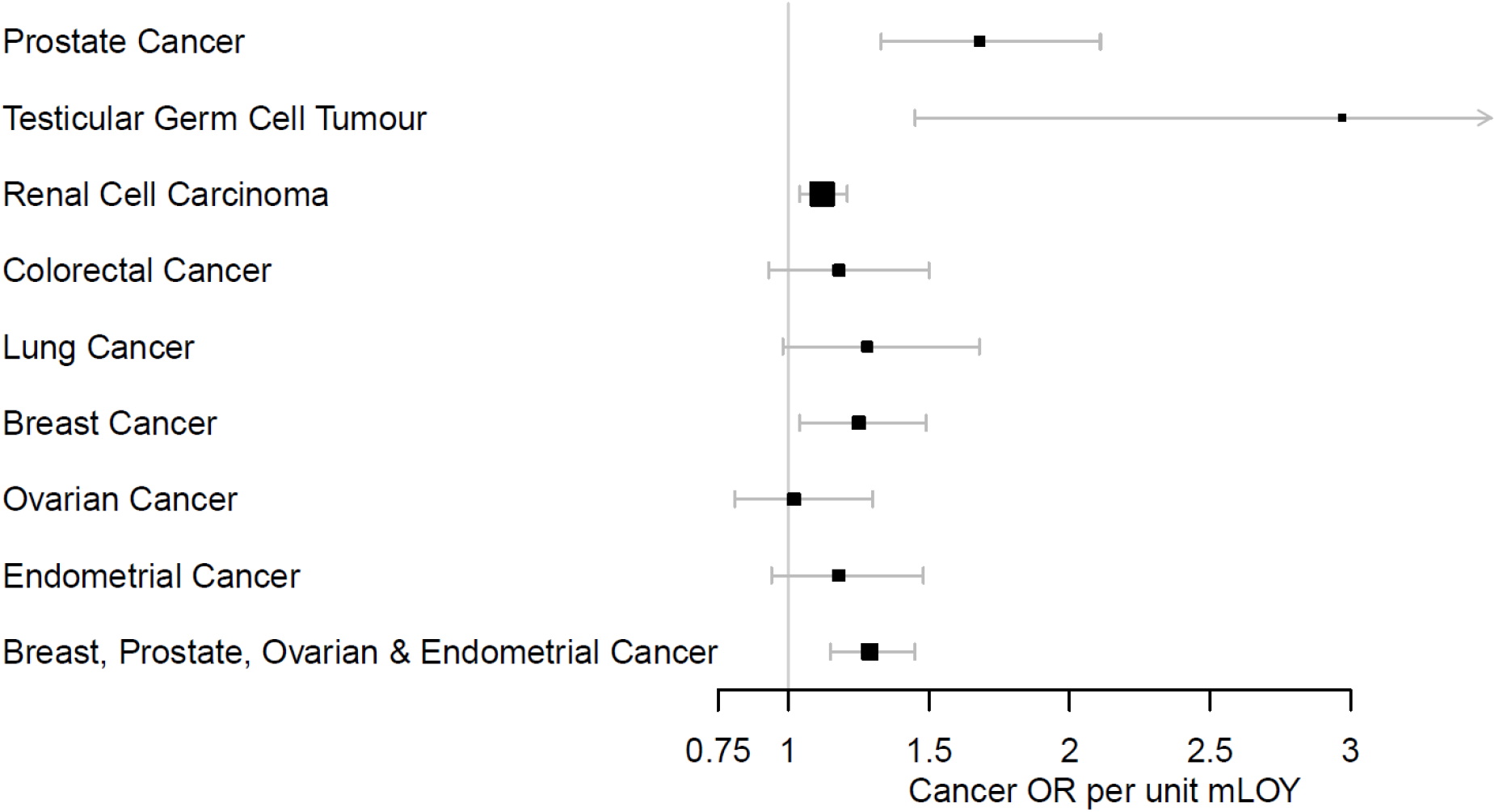
Association between a genetic risk score of the 156 LOY-associated variants and cancer.

### Genetic predisposition to LOY is associated with health outcomes in women

Mosaic LOY in blood cells has been associated with a broad range of diseases, which if causal is likely explained by one (or both) of two mechanisms: either LOY in leukocytes has a direct physiological effect, for example through impaired immune function, and/or it acts as a barometer and readily detectable manifestation of genomic instability occurring in parallel in other tissues. Ideally, this question would be addressed by assessing clonal mosaicism in large population studies where DNA was extracted from a broad range of cell and tissue types. In the absence of such a study, we hypothesised that testing the relevance of our identified LOY associated variants in women would help inform this – any association between the two could not be explained by a direct effect of LOY, given that females are XX.

To assess this we tested a polygenic risk score comprised of our 156 lead variants for association with three female-specific cancers – breast, endometrial and ovarian (Figure 4, **Table S15**). We observed a significant association with breast cancer (OR 1.25 (1.04-1.49) P=0.016) and directionally consistent results in the smaller endometrial (OR 1.18 (0.94-1.48), P=0.14) and ovarian (OR 1.02 (0.81-1.30), P=0.86) studies.

We next tested the same score on a female-specific non-cancer trait also underpinned by genome instability – age at natural menopause. Previous human and animal studies have shown menopause age is substantially biologically determined by the ability of oocytes to detect, repair and respond to DNA damage^29,30^. We found that genetically increased risk of LOY was associated with later age at menopause (P=0.003, **Table S16**), with the *CHEK2* locus individually reaching genome-wide significance (P=7.9×10^−22^). A repeated genetic risk score analysis excluding *CHEK2* retained significance (P=0.017).

Given this observation that genetic susceptibility to LOY in leukocytes is impacting broader biological systems in these women, it is reasonable to speculate that actual LOY in leukocytes in men similarly represents a biomarker of genome instability in other cell and tissue types.

### Exploring the impact of LOY at the level of a single cell

To help understand if and why LOY may provide a growth advantage to a cell, and the potential mechanisms linking LOY to disease, we performed single cell transcriptomic analyses (scRNAseq) using the 10X Genomics Chromium Single Cell 3’ platform. This was performed on peripheral blood mononuclear cells (PBMCs) collected from 19 male donors (aged 64-89), unselected for any measure of clonal mosaicism. After standard quality control steps (see methods), we sequenced and profiled gene expression across 86,160 single cells. Under normal conditions, blood cells express a set of genes located in the male specific region of the Y chromosome (MSY). The LOY status of individual cells could therefore be determined by the absence of expression from these genes, which we identified in 13,418 of the cells (15.6% across all cells, ranging from 7-61% within individuals).

Across the autosomal genome the most strongly differentially expressed gene between cells with and without the Y chromosome was *TCL1A* (Figure 5). This gene maps to one of our identified genetic variants (rs2887399, 162bp away), where the LOY risk increasing allele is associated with higher *TCL1A* expression in blood (**Table S10**). The single cell data showed that, among the major types of leukocytes, the *TCL1A* gene was expressed only in B-lymphocytes (Figure 5) and LOY was detected in 11.3% of these cells, ranging from 2% to 56% within individuals. B-lymphocytes without the Y chromosome (cell N=277) had 75% higher normalized *TCL1A* expression compared to those with a Y chromosome (N=2,459, Wilcoxon test in Seurat: fold change=1.75, P<0.0001). We also performed an in-house resampling test to evaluate this difference and validated a substantial upregulation of *TCL1A* in LOY cells (resampling test: fold change=1.68, P<0.0001) (Figure 5). An analysis within each individual demonstrated single cells with LOY had consistently higher *TCL1A* expression, ruling out any bias by *TCL1A* genotype (**Figure S3**).

**Figure 5.**
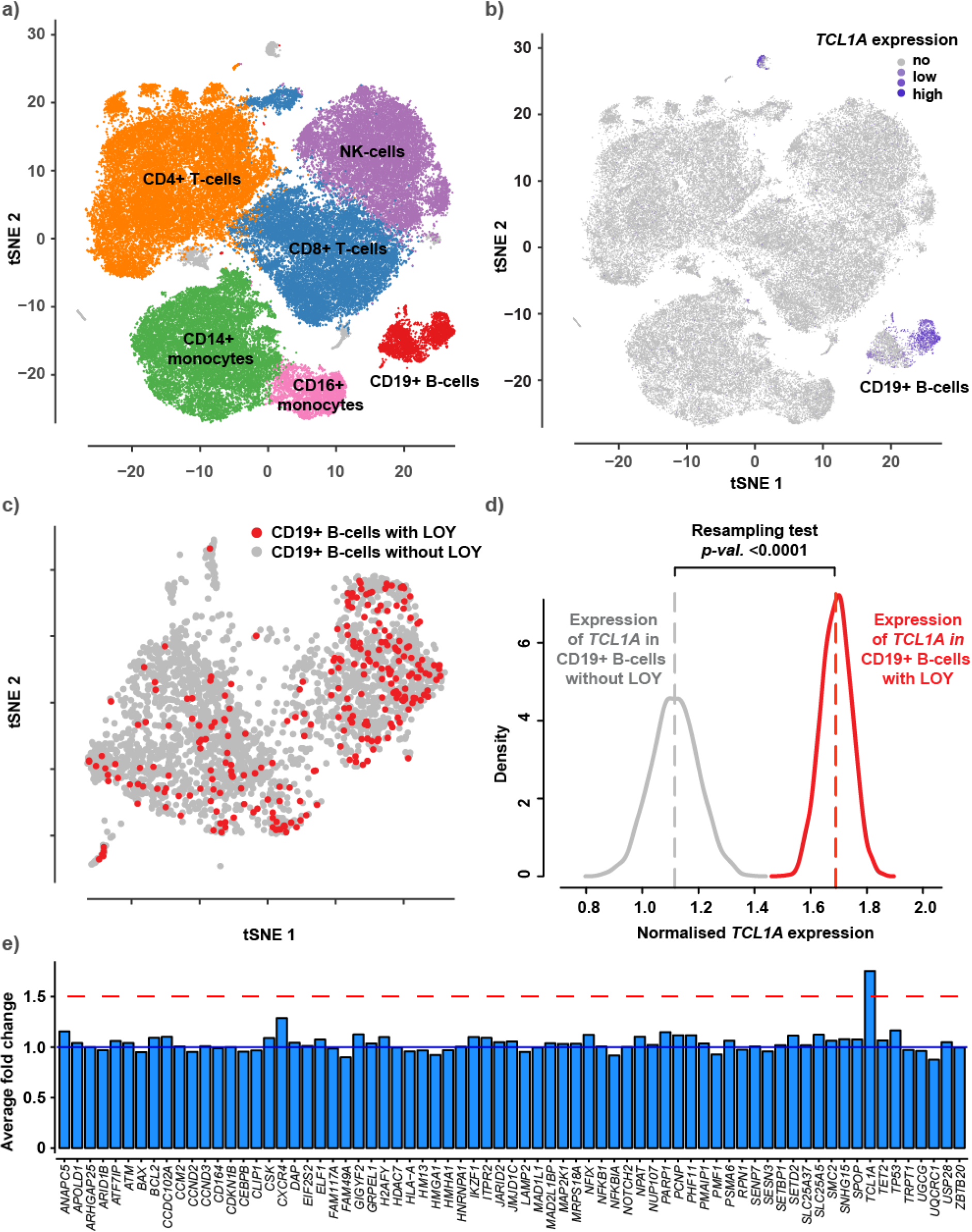
Single-cell RNA sequencing results. Panel a shows clustering and identification of cell types using a tSNE plot generated from a pooled dataset including 86160 PBMC’s isolated from peripheral blood samples collected from 19 male donors. The *TCL1A* gene was expressed in the B-lymphocytes as indicated by blue color in panel **b.** Analysis of LOY status in the B-lymphocytes identified 277 cells with LOY, plotted in red color in panel **c.** Panel **d** display the result from a resampling test performed to compare the expression of *TCL1A* in LOY B-lymphocytes with its expression in non-LOY B-lymphocytes. The grey and red curves in panel **d** represent the resampled distribution of *TCL1A* expression in non-LOY and LOY cells, respectively. The resampling test established an increased expression of *TCL1A* in B-lymphocytes with LOY (fold change=1.68, *p*<0.0001). Panel **e** display fold changes in gene expression between LOY and non-LOY B-lymphocytes for 71 selected genes from the list of genes mapping to the 156 index variants. Genes expressed in >5% of the investigated B-lymphocytes were included. The blue line at fold change 1 in panel e represents no differential expression and the red line shows the level of 50% overexpression in LOY cells.

To evaluate the magnitude of the 75% overexpression of the *TCL1A* gene in LOY B-lymphocytes, we compared the expression changes of other genes proximal to our identified GWAS loci. Of the genes we prioritized at each of our GWAS loci (“consensus genes”, **Table S8**), 71 were expressed in >5% of the B-lymphocytes and included in the comparison, but only *TCL1A* demonstrated significant fold change (Figure 5).

These data provide a possible explanation for the growth advantage conferred to cells missing a Y chromosome. *TCL1A* is a known oncogene, the product of which (TCL1) mediates intracellular signalling and stimulates cell proliferation and survival^31^. TCL1 has also been associated with down regulation of p53 activity through activation of *MDM2*^32^. While p53 action is associated with the G1/S cell cycle checkpoint, it also has post-mitotic checkpoint functions^33^. It could therefore be hypothesised that LOY causes down regulation of p53 via *TCL1A* upregulation, leading to enhanced proliferation and inhibited apoptosis of LOY cells. The independent effect of *TCL1A* genotype also suggests a possible bidirectional involvement for *TCL1A*. Ultimately further experimental work will be required to fully elucidate the aetiological implications of altered *TCL1A* expression in these cells.

## Discussion

This study provides several advances in our understanding of the likely underlying biology and probable consequences of mosaic Y chromosome loss in circulating leukocytes. Our newly developed detection algorithm (PAR-LOY) and expanded sample size led to an 8-fold increase in the number of associated genetic determinants, which we use to make several important observations.

The origin of LOY at the level of a single cell is perhaps most readily explained by chromosome mis-segregation events during mitosis. Consistent with this, many of the identified loci harbour nearby genes involved in key mitotic processes (Figure 6), notably central components of condensin which affects mitotic chromosome structure (*NCAPG2, SMC2*)^34^, and the assembly, structure and function of the kinetochore (*CENPN, CENPU, PMF1, ZWILCH*) and spindle (*SPDL1*), which together form the main machinery of chromosome congression and segregation^35,36^. *MAD2L1* (alongside *MAD1L1* and *MAD2L1BP*) and *ZWILCH* are core components of the mitotic spindle assembly checkpoint^37^, which ensures that chromatids are bi-orientated at the metaphase plate and under bipolar tension before disinhibiting the anaphase-promoting complex (of which *ANAPC5* is a component) to allow progression from metaphase. Many genes governing wider cell cycle progression, including cyclins (*CCND2, CCND3*), regulators of cyclin (*CDKN1B, CDKN1C, CDK5RAP1*) and major checkpoint kinases (*ATM*) are also identified here, emphasising the importance of processes across the cell cycle in determining LOY. A remainder of the genes that we identify encode proteins involved in sensing and responding to DNA damage (*SETD2, DDB2, PARP1, ATM, TP53, CHEK2*) and apoptotic processes (*PMAIP1, SPOP, LTBR, SGMS1, TP53INP1, DAP)*. Of note, *FANCL* – the nearest gene to one of our lead variants – is vital to DNA interstrand crosslink repair and mutations in this gene have been linked to a rare autosomal form of Fanconi Anaemia, characterised by cytogenetic instability and chromosome breakage^38^. The Bcl-2 family, a conserved set of proteins that regulate caspase-mediated apoptosis by controlling mitochondrial release of Cytochrome-C, are also particularly well-represented (*BCL2, BAX, BCL2L1, BCL2L11*)^39^. These themes are consistent with the hypothesis that, secondary to the initial mis-segregation event, clonal expansion of LOY cells requires an environment permissive to proliferation of aneuploid cells, in which normal processes to detect and terminate these cells are avoided.

**Figure 6.**
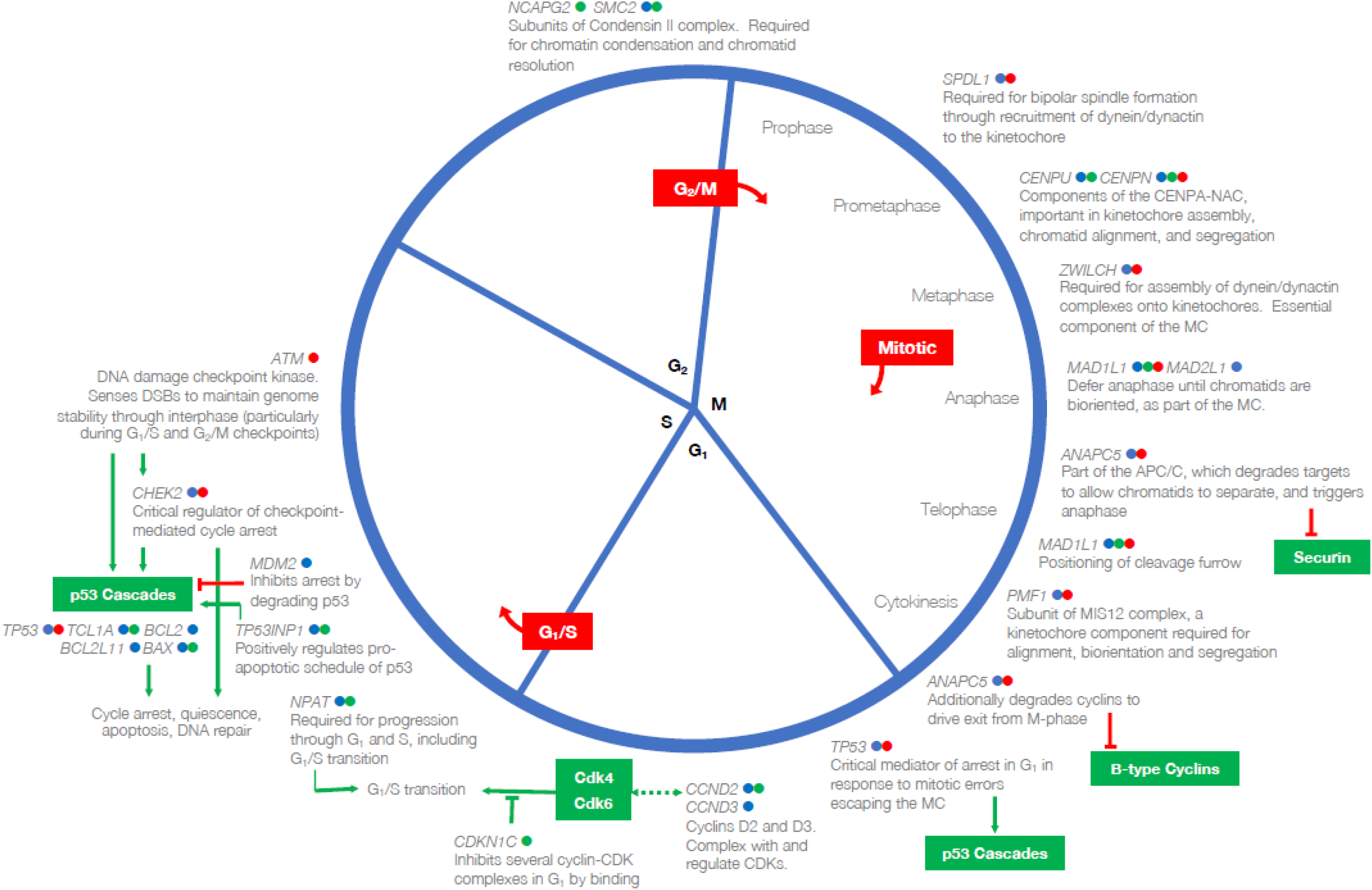
Many LOY-associated genes converge on mechanistic and regulatory aspects of the cell cycle. All genes shown have been prioritized as potentially functional genes at our reported GWAS loci; gene symbols may be shown more than once. Coloured indicators next to each gene symbol specify the type of evidence on which it has been prioritized at its respective locus: blue, nearest protein-coding gene; green, eQTL; red, contains a highly correlated non-synonymous variant. Red boxes indicate each of the three known cell cycle checkpoints. Red inhibition connectors denote that a target is inhibited by degradation, green by binding. Green arrows indicate a signaling cascade and its effector or final physiological effect. Bidirectional dashed green arrows indicate the formation of a complex between the products of the two connected genes. Excepting p53, proteins contained within green boxes have not been implicated in this GWAS, but are important interactors of implicated genes. CENPA-NAC, CENPA nucleosome-associated complex; APC/C, anaphase-promoting complex/cyclosome; MC, mitotic checkpoint; CDK, cyclin-dependent kinase.

A link between LOY and cancer susceptibility seems plausible conceptually, given the nature of the genes identified. Here, we find substantial overlap of LOY associated variants across known cancer susceptibility loci, somatic drivers of tumour growth and genes targeted by licensed or in-trial cancer therapeutics. A notable example is the target of PARP inhibitors *PARP1*, where the lead SNP is highly correlated with a missense variant (V762A), the minor allele for which (the alanine substitution) is protective for LOY and has experimentally been shown to reduce PARP-1 catalytic activity by 30-40%^40^. More broadly, we found evidence for a systematic relationship between genetic susceptibility for LOY and risk of breast, prostate, testicular and renal cell carcinomas (Figure 4).

Based on our observations, we propose that LOY is determined by a “common soil” of shared mechanisms that predispose to genome instability and cancer. This is perhaps most readily apparent with the observation that genetic susceptibility to LOY is associated with cancer susceptibility in women and age at natural menopause. Although in aggregate we found that LOY associated variants tended to delay menopause age, there was substantial heterogeneity in dose-response. This was exemplified by individually significant associations for LOY risk increasing alleles with both earlier (mapped genes *MON1A, NR6A1, FBXL20*) and later menopause (*PMF1, JMJD1C, USP35/GAB2, APOLD1/CDKN1B, PHF11/RCBTB1, HEATR3, TP53* and *CHEK2)*. We hypothesised that the direction of effect a LOY associated risk allele has on menopause may shed light on its mechanism of effect. Genetic determinants of LOY must broadly act either by promoting chromosomal instability or facilitating the clonal expansion of aneuploid cells, both of which are potentially cancer causing mechanisms. An allele that promotes clonal mosaicism through predisposition to chromosome mis-segregation or repair/generation of DNA damage would likely be associated with an earlier menopause due to acquired DNA damage in oocytes and their subsequent elimination. This principle is most evident in mice and humans with *BRCA1/2* loss of function, where diminished double strand break repair in oocytes triggers apoptosis and depletion of the ovarian reserve^30,41,42^. In contrast, any process that impairs DNA damage sensing or programmed cell death may also promote clonal mosaicism (via greater tolerance of damaged cells) but lead to a *later* menopause. The clearest example of this is *CHEK2*, which in mice is essential for culling oocytes bearing unrepaired DNA double-strand breaks^43^. Reduced activity of this gene would therefore lead to the survival of defective oocytes (hence later menopause), supported for the first time in humans by our observation that *CHEK2* loss of function is associated with later age at menopause in women and increased LOY in men. The overall trend for LOY associated loci to be associated with delayed menopause suggests that many genetic determinants may act to inhibit apoptosis and the elimination of defective leukocytes. Further experimental work in animal and cellular models should aim to investigate more thoroughly the mechanisms linking each of our putatively highlighted genes to clonal mosaicism and broader outcomes.

We also note overlap between our identified LOY associated loci and other complex traits and diseases. For example, seven of our current LOY signals are correlated with previously reported^44^ susceptibility loci for Type 2 diabetes (*TP53INP1, SUGP1, KCNQ1, CCND2, EIF2S2, PTH1R* and *BCL2L11*). At six of these overlapping loci, the LOY risk-increasing allele also increases the risk of Type 2 diabetes. *CCND2* encodes Cyclin D2, the major D-type cyclin expressed in pancreatic β-cells and is essential for adult β-cell growth^45^. *TP53INP1* is a p53-inducible gene, whose product regulates p53-dependent apoptosis. Additionally, the LOY-associated genes encoding cyclins and cyclin-dependent kinases, *CCND3, CDKN1B* and *CDKN1C*, are also implicated in pancreatic β-cell growth and maturation. We hypothesise that the previously reported association between clonal mosaicism in blood and T2D^15,46^ may reflect a common susceptibility to cell cycle dysregulation and genome instability, which lead to both increased clonal mosaicism and reduced pancreatic β-cell mass.

Finally, the “common soil” hypothesis discussed above does not preclude the possibility that LOY in leukocytes also has a direct role in disease, for example through impaired immune function^47^. A growing awareness of the physiological importance of chromosome Y outside of reproductive development challenges the view of this chromosome as a “genetic wasteland”^48^. The male-specific region (MSY) encodes 27 distinct proteins, with roles in fundamental processes such as chromatin modification (*KDM5D, UTY*), gene transcription (*ZFY*) and translation (*DDX3Y, EIF1AY* and *RPS4Y1*). Indeed, our observation in single-cell RNA sequencing data that leukocytes with LOY have dysregulated autosomal gene expression supports the notion of a direct physiological effect.

We hope that future experimental studies may build on these observations, yielding further insights into mechanisms that may have broad relevance to a range of cancers and other ageing-related diseases.

## Acknowledgements

This research has been conducted using the UK Biobank Resource under application 9905. This work was supported by the Medical Research Council [Unit Programme number MC_UU_12015/2]. Full study-specific and individual acknowledgements can be found in the supplementary information.

## Methods

### Phenotype preparation in UK Biobank

We developed a new statistical approach for identifying male individuals with LOY based on allele-specific genotyping intensities in the pseudoautosomal region (PAR) of the sex chromosomes. In contrast to previous work that has quantified Y chromosome loss based on median genotyping intensity over the non-pseudoautosomal region of the Y chromosome (mLRR-Y)^12–15^, our approach leverages the diploid nature of the PAR to ascertain mosaic Y loss based on differences between maternal (X PAR) vs. paternal (Y PAR) allelic intensities at heterozygous sites: mosaic Y loss causes Y PAR intensities to decrease relative to X PAR intensities. This intuition can be harnessed even in population cohorts in which absolute phase information (i.e., information about maternal vs. paternal inheritance of alleles) is unavailable: we can overcome this obstacle by performing statistical phasing and subsequently identifying evidence of an imbalance in allelic intensities between the two statistically phased haplotypes (accounting for the possibility of phase switch errors)^5,6^. In general, the signal produced by phased allelic imbalances is typically much cleaner than estimates of total genotyping intensities (e.g., mLRR-Y), as the latter can vary substantially across the genome due to technical artefacts^49^.

We applied this approach to blood DNA genotyping intensity data from the full UK Biobank cohort (described extensively elsewhere^50^, analyzing 1,239 genotyped variants on PAR1 that passed QC (out of 1,301 total PAR1 variants). (We ignored the much-shorter PAR2, which only contained 56 genotyped variants, of which 37 passed QC.) To maximize phasing accuracy, we phased the full cohort including both males and females using Eagle2^51^, after which we restricted our attention to males. We called mosaic chromosomal alterations (mCAs) in PAR1 using a slightly modified version of the pipeline we described previously^6^. Specifically, in our hidden Markov model, we increased the probability of starting in a mosaic state to 0.2 (reflecting our call set; see below), and we also postprocessed our PAR1 mCA calls to identify likely mosaic Y loss events based on two criteria: (i) mCA spans the full PAR1 region; and (ii) observed mean log_2_ R ratio (LRR) is more consistent with a mosaic loss event than a CNN-LOH or gain (after taking into account the s.e.m. of LRR and an empirical prior on mCA copy numbers^6^. This procedure produced 44,709 mCA calls in PAR1 (at an estimated false discovery rate of 0.05) among 220,924 males passing sample QC, of which 43,306 were classified as likely LOY. These calls contained an average of 321 heterozygous variants on PAR1 passing QC that were usually phased perfectly (no switch errors detected by the hidden Markov model in 72% of calls).

Recalled age at natural menopause (ANM) was available in 106,237 women with genetic data. We included women with ANM who were 40–60 years of age in our analyses, excluding those with menopause induced by hysterectomy, bilateral ovariectomy, radiation or chemotherapy and those using hormone replacement therapy (HRT) before menopause.

### Genetic association testing in UK Biobank

We used genetic data from the “v3” release of UK biobank^50^, containing the full set of HRC and 1000G imputed variants. In addition to the quality control metrics performed centrally by UK Biobank, we defined a subset of “white European” ancestry samples using a K-means clustering approach applied to the first four principle components calculated from genome-wide SNP genotypes. Individuals clustered into this group who self-identified by questionnaire as being of an ancestry other than white European were excluded. After application of QC criteria, a maximum of 205,011 male participants were available for analysis with genotype and phenotype data. Association testing was performed using a linear mixed models implemented in BOLT-LMM^52^ to account for cryptic population structure and relatedness. Only autosomal genetic variants which were common (MAF>1%), passed QC in all 106 batches and were present on both genotyping arrays were included in the genetic relationship matrix (GRM). Genotyping chip, age at baseline and 10 genetically derived principal components were included as covariates.

We defined statistically independent signals (described as lead or index variants) using 1Mb distanced-based clumping across all imputed variants with P<5×10^−8^, an imputation quality score > 0.5 and MAF > 0.1%. Genome-wide significant lead variants that shared any correlation with each other due to long range linkage disequilibrium (r^2^>0.05) were excluded from further consideration. These loci were additionally augmented using approximate conditional analyses implemented in GCTA^53^. Here, secondary signals were only considered if they were uncorrelated (r^2^<0.05) with a previously identified index variant and genome-wide significant pre and post conditional analysis.

The total trait variance of all genotyped SNPs was calculated genome-wide and per-chromosome using restricted estimate maximum likelihood (REML) implemented in BOLT-LMM^52^. The corresponding observed-scale estimate was transformed to the liability-scale^54^.

### Replication

Replication was performed in three independent studies using two separate techniques.

Firstly, we used data generated from the customer base of 23andMe Inc, a consumer genetics company. Genotyping array quality control, imputation and downstream association testing for this study has been described extensively elsewhere^55^. All individuals provided informed consent and answered surveys online according to 23andMe’s human subjects protocol, which was reviewed and approved by Ethical & Independent Review Services, a private institutional review board (http://www.eandireview.com). DNA extraction and genotyping were performed on saliva samples by National Genetics Institute (NGI), a CLIA licensed clinical laboratory and a subsidiary of Laboratory Corporation of America Mosaic LOY was estimated by calculating the mean log-R ratio (normalised signal intensity) across 274 SNPs on the male-specific region of the Y chromosome that are shared and perform well across genotyping platforms, using the protocol described previously^14^. Imputation was performed using a combination of the May 2015 release of the 1000 Genomes Phase 3 haplotypes^56^ with the UK10K imputation reference panel^57^. Genetic association testing was performed using linear regression in 653,019 male research participants of European ancestry, using age, genetically derived principal components and genotyping platform as covariates. Results were adjusted for a genomic control inflation factor of 1.129.

Secondly, we analyzed whole-blood genome sequences of 8,715 Icelandic males^58^ (age range 41-105 years, mean 63 years), that had been whole-genome sequenced by Illumina method to a mean depth of 37x. As an estimate of chromosome Y copy-number we used the average read depth over chromosome Y, using exclusively X-degenerate regions. This was computed by samtools from bam files aligned to hg38 and normalized by genome-wide sequencing coverage for the subject. A total of 12 outlier individuals (copy-number greater than 1.25) were excluded. Association analysis was performed using BOLT-LMM^52^ after inverse normal transformation and adjustment for age at bleeding. Effect sizes for log2(chrY copy-number) were estimated using robust linear regression (rlm from R package MASS).

Third, we used a sample of 95,380 Japanese ancestry men from the BBJ project, a study which has been described extensively elsewhere^59^. The study was approved by the ethical committees in the Institute of Medical Science, the University of Tokyo and RIKEN Center for Integrative Medical Science. Mosaic LOY in blood was estimated using the quantitative mLRR-Y measure, using a similar protocol as previous studies^14^. Association testing was performed using a linear mixed model implemented in BOLT-LMM^52^, including age, smoking, disease status and chip array as a covariate.

### Genomic feature enrichment

We used a previously modified version of GoShifter^26,60^ to calculate the enrichment of fine-mapped (PP ≥ 0.10) and not fine-mapped (PP < 0.10) variants with genomic annotations by locally shifting the annotations and computing overlaps to approximate the null distribution. Z-scores and odds ratios were calculated from 1000 permutations, and typical two-tailed p-values are calculated from the z-score statistic. All annotations were obtained as described previously^26^.

In order to identify which tissue types were most relevant to genes involved in LOY, we applied LD score regression^61^ to specifically expressed genes (“LDSC-SEG”)^62^ and g-chromVAR to hematopoietic accessible chromatin^26^. For LDSC-SEG, cell-type specific analyses using GTEx and Epigenome Roadmap annotations were performed using the data available on the LDSC-SEG resource page (https://github.com/bulik/ldsc/wiki/Cell-type-specific-analyses). For g-chromVAR, hematopoietic specific analyses were performed using ATAC-seq count matrices as previously processed (https://github.com/caleblareau/singlecell_bloodtraits). g-chromVAR estimates were averaged across 10 different random background peak sets. We note that, similar to the derivation of cell-type specific features or SEGs in LDSC, g-chromVAR z-scores represent relative enrichment for specific cell-types compared to other input cell-types, which allows for discrimination between closely related cell types in the hematopoietic lineage.

### Gene expression integration

We used two approaches to map associated genetic variants to genes via expression effects (eQTLs) in whole blood. Firstly, Summary Mendelian Randomization (SMR) uses summary-level gene expression data to map potentially functional genes to trait-associated SNPs^63^. We ran this approach using a meta-analysis of whole blood eQTL data from 31,684 individuals^64^. Only transcripts with no evidence of pleiotropic effects, as assessed by the HEIDI metric were considered^63^. Secondly, we used the recently described Transcriptome-wide Association Study (TWAS) approach^65^ to infer gene expression association using three whole blood datasets (Young Finns Study, Netherlands Twin Registry cohorts and GTEx v6). All data used is available here: http://gusevlab.org/projects/fusion/. For all analyses significance thresholds were set to adjust for the number of tested performed.

### Pathway enrichment analysis

Pathway analysis was performed using two distinct approaches – STRING^66^ and MAGENTA^67^. For STRING, only the gene closest to one of the 156 lead index variants was included in the analysis. In contrast, MAGENTA performs enrichment analysis using the full genome-wide summary statistic data.

We used the Open Targets Platform (https://www.targetvalidation.org/) to define gene sets comprising genes involved in cancer susceptibility (N=249), somatic drivers of tumour growth (N=394), targets of approved or in trial cancer therapies (N=458), “affected pathways” (N=216 with score = 1) and finally an overall aggregated score for involvement in cancer (N=934 with score = 1). The various data sources, and approach applied by Open Targets to score and prioritise target genes within each of these categories is described in full at https://docs.targetvalidation.org/getting-started/scoring). We arbitrarily defined gene set membership based on an assigned score > 0.8 unless otherwise specified. These pathways were tested for enrichment in downstream analyses using MAGENTA.

### Fine-mapping

Regions for fine-mapping were defined by extended 0.5 Mb in both directions from each sentinel and merging when regions overlapped, resulting in 126 total regions. All variants in these regions with MAF > 0.005 and INFO > 0.6 were fine-mapped. Dosage LD was estimated from the UKB genotype probability files (.bgen) using 167,020 unrelated white British male individuals (http://www.nealelab.is/uk-biobank/). Fine-mapping was them performed using v1.3 of the FINEMAP software^68^ with default settings allowing for up to 5 causal variants in each region. The UCSC genome browser was used to view individual variants along with hosted features^69^.

### Integration with cancer data and modelling LOY as a causal exposure

The NHGRI-EBI GWAS Catalogue database was accessed and downloaded on June 25, 2018. The downloaded file was curated to only include studies in which cancer is the associated disease and further filtered to remove variants with association p-values greater than 5 × 10^−8^. Due to a potential lag between the time a new GWAS is published and included in the NHGRI-EBI GWAS Catalogue, a supplementary literature search of PubMed was performed to identify additional reports of cancer susceptibility studies that were not included in the GWAS Catalogue. The literature search was completed on July 18, 2018. LDlink^70^ was used to identify published cancer GWAS-associated genetic variants which are in linkage disequilibrium (LD) (r^2^≥0.1 based on the 1000 Genomes Project European Population data) with one of the 154 LOY lead SNPs. Associations with haematological malignancies were excluded and additional associations were identified by manual searches.

The relationship between LOY-associated variants and cancer was assessed using a two-sample summary statistic based Mendelian randomisation analysis. Linear regressions of the cancer log odds ratios (logOR) for each available SNP on the LOY beta coefficients were carried out, weighted by the inverse of the variance of the cancer logORs. This is equivalent to an inverse-variance weighted meta-analysis of the variant-specific causal estimates^71^. Because of evidence of over-dispersion (i.e. heterogeneity in the variant-specific causal estimates), the residual standard error was estimated, making this equivalent to a random-effects meta-analysis. Unbalanced horizontal pleiotropy was tested based on the significance of the intercept term in MR-Egger regression^72^.

Summary statistics for the association between the genetic variants and risk of prostate cancer were obtained from the PRACTICAL/ELLIPSE consortium, based on GWAS analyses of 67,158 prostate cancer cases and 48,350 controls^73^. Testicular cancer summary statistics were obtained from two GWAS studies conducted at the Institute of Cancer Research comprising 4,192 testicular cancer cases and 12,368 controls^74,75^. The renal cancer analysis used summary statistics from the Kidney Cancer GWAS Meta-Analysis Project of 10,784 cases of renal cell carcinoma and 20,406 controls^76^. Colorectal cancer summary statistics were from eight UK-based GWAS studies, totalling 22,372 colorectal cancer cases and 44,271 controls^77,78^. The summary statistics for overall lung cancer were from GWAS analyses of 29,266 lung cancer cases and 56,450 controls conducted by the International Lung Cancer Consortium^79^. The breast cancer analysis was based on summary statistics from GWAS analyses of 105,974 breast cancer cases and 122,977 controls conducted by the Breast Cancer Association Consortium (BCAC)^80^, including summary statistics from analyses restricted to cases with estrogen receptor positive or estrogen receptor negative breast cancer^81^. Summary statistics for the ovarian cancer analysis were from GWAS studies of 25,509 ovarian cancer cases and 48,941 controls conducted by the Ovarian Cancer Association Consortium (OCAC)^82^. The endometrial cancer results were from GWAS studies of 12,906 endometrial cancer cases and 108,979 controls from the Endometrial Cancer Association Consortium (ECAC)^83^. In addition, MRs for breast and ovarian cancer risk specifically in carriers of a *BRCA1* or a *BRCA2* mutation were carried out using results from GWAS studies conducted by the CIMBA consortium^81,82^. There was some overlap in the control subjects used by the breast, ovarian, endometrial and colorectal cancer studies, and between the control subjects used in the prostate cancer study, the colorectal cancer study and one of the testicular cancer studies. All the cancer MRs were based on summary statistics from analyses restricted to participants with European ancestry.

The pan-cancer summary statistics (breast, prostate, ovarian, and endometrial) were derived using a three step procedure. First, the tetrachoric correlation of binary transformed Z-scores was used to estimate the correlation between individual-cancer summary statistics that is attributable to control sample overlap^84^. Second, individual-cancer summary statistic standard errors were decoupled to account for the estimated correlation^85^ and third, the METASOFT software^86^ was used to perform fixed effect inverse-variance weighted meta-analyses for the combination of four cancers.

### Sample preparation for single cell gene expression study

Blood samples from 19 elderly men (median age=80, range=64-89) admitted to the Geriatrics Department at Uppsala University Hospital (Uppsala, Sweden) were collected in BD Vacutainer CPT cell separation tubes containing sodium citrate and stored on ice. The PBMC fraction was isolated from the whole blood samples by density gradient centrifugation following manufacturer instructions (Becton, Dickinson and Company, Franklin Lakes). PBMCs were collected and suspended in cold 1X PBS solution with 0.04% BSA. Cell concentrations were measured using an EVE cell counter (NanoEnTek, Seoul) and diluted to a concentration of 10^6^ cells/ml. All the prepared samples had a cell viability above 90%. The local research ethics committee approved the study and all participants provided their informed consent.

### Single cell workflow

We performed single cell RNA sequencing (scRNAseq) using the 10X Chromium Single Cell 3’ gene expression solution (10X Genomics, Inc.) at the SNP&SEQ Technology Platform at Uppsala University (Sweden). This scRNAseq technology is based on gel beads loaded with barcoded oligos mixed with single cells and enzymes, before captured in droplets (GEMs). The transcripts present in individual cells are barcoded with UMI’s (unique molecular identifiers) and used to prepare standard sequencing libraries. All transcripts from single cells get barcoded with the same index sequence allowing for the transcripts from thousands of single cells to be pooled together in a single sequencing run and allowing transcriptional profiling of individual cells. The barcoding and library construction were performed for the 19 PBMC samples using the Chromium Single Cell 3’ Reagent kit (cat# 120236/37/62) according to the manufacturer protocol (CG00052 Single Cell 3’ Reagent Kit v2 User Guide). The entire procedure, from blood sampling to construction of GEM’s was accomplished within 5 hours. The generated single cell libraries were sequenced using a NovaSeq 6000 instrument (Illumina, Inc., San Diego) at the SNP&SEQ Technology Platform and generated a median of 64900 reads per cell (range=35213-111643).

### Single-cell bioinformatics pipeline

Sequenced reads were mapped to the human reference (GRCh37/hg19) using the software Cellranger v 2.0.2 (10X Genomics, Inc.). Cellranger produces a count-matrix for each experiment containing the UMI barcodes using sequence information from the 3’ end of each transcript in every single cell. We used the R library Seurat (v2.3.1) for further processing and implemented the standard Seurat workflow7. Specifically, standard QC-steps were performed including removal of apoptotic cells (i.e. cells with a large fraction of mitochondrial RNA) as well as removing cells with low sequencing coverage and/or a low number of expressed genes, as recommended. Following QC-steps, normalization of the gene expression within each single cell was performed using the function “NormalizeData”. Genes with the most differential expression within single cells were identified for cluster analysis and normalized gene expression was scaled using the “ScaleData” function. Principal components were calculated using the most variable genes and the number of significant principal components was determined. Clustering of the dataset was performed using the function “FindClusters” and cell types for each cluster were determined using canonical marker genes. Refined clustering was achieved by reclustering within the identified cell types using the above pipeline on subsets of the data. The tSNE plots were produced using the generated principal components.

### Determination of LOY in single cells

The LOY status for each sequenced cell was determined under the assumption that cells with LOY would not express genes located in the male specific part of chromosome Y (MSY). Hence, non-LOY blood cells are normally expressing a series of genes located in the MSY: DDX3Y, EIF1AY, KDM5D, RPS4Y1, USP9Y, UTY and ZFY. We took advantage of this information and thus scored LOY in cells without any transcripts from the genes located in the MSY. Only cells with expression and good quality data from genes located on the autosomal chromosomes were included.

### Single-cell statistical analyses

To compare differences in autosomal gene expression between LOY cells and non-LOY cells we first performed WilcoxDETest’s, implemented in the R library Seurat (v2.3.1). We also developed an inhouse random sampling algorithm to compare the gene expression in LOY cells with non-LOY cells within specific cell types. First, we established the observed gene expression in LOY cells in the cell type under investigation, by calculating the mean normalized expression values in all subjects, within all LOY cells of the tested cell type. Next, we randomly selected from all subjects, a number of cells from the non-LOY cells of the examined cell type, and calculated the mean normalized expression in the resampled cells. To avoid biases caused by inter-individual variation, we programmed the sampling algorithm to sample an equal number of non-LOY cells as observed LOY cells within each subject. For example, from subjects with 100 LOY cells of a specific cell type, the same number of non-LOY cells from the same cell type was sampled from the set of non-LOY cells. The resampling of non-LOY cells from all subjects was repeated 50.000 times and for each iteration, the mean normalized expression of the investigated gene in the resampled cells was calculated. The resampled data represents a weighted expression level of the examined gene in non-LOY cells within specific cell types and thus, the resampled distribution represents the normalized expression of the investigated gene in non-LOY cells. The range of variation of gene expression in LOY cells was estimated in a similar fashion, by resampling of a subset of the LOY cells within each subject. Exact p-values were calculated by comparing the observed mean expression in LOY cells to the resampled distribution of non-LOY cells. All statistical analyses were performed using R v. 3.4.4.

