## Supplementary Information and Figures for "Genetic predisposition to mosaic Y chromosome loss in blood is associated with genomic instability in other tissues and susceptibility to non-haematological cancers"

**Supplementary text and figures**

1. Supplementary figures

   S1 - Liability-scale heritability explained vs chromosome size.

   S2 - Cell and tissue type enrichment estimated using LDSC-SEG

   S3 - Differential expression of the *TCL1A* gene in B-lymphocytes with and without the Y chromosome within individual subjects

2. Consortium authorship

3. Acknowledgements

**Figure S1 | Liability-scale heritability explained vs chromosome size.**

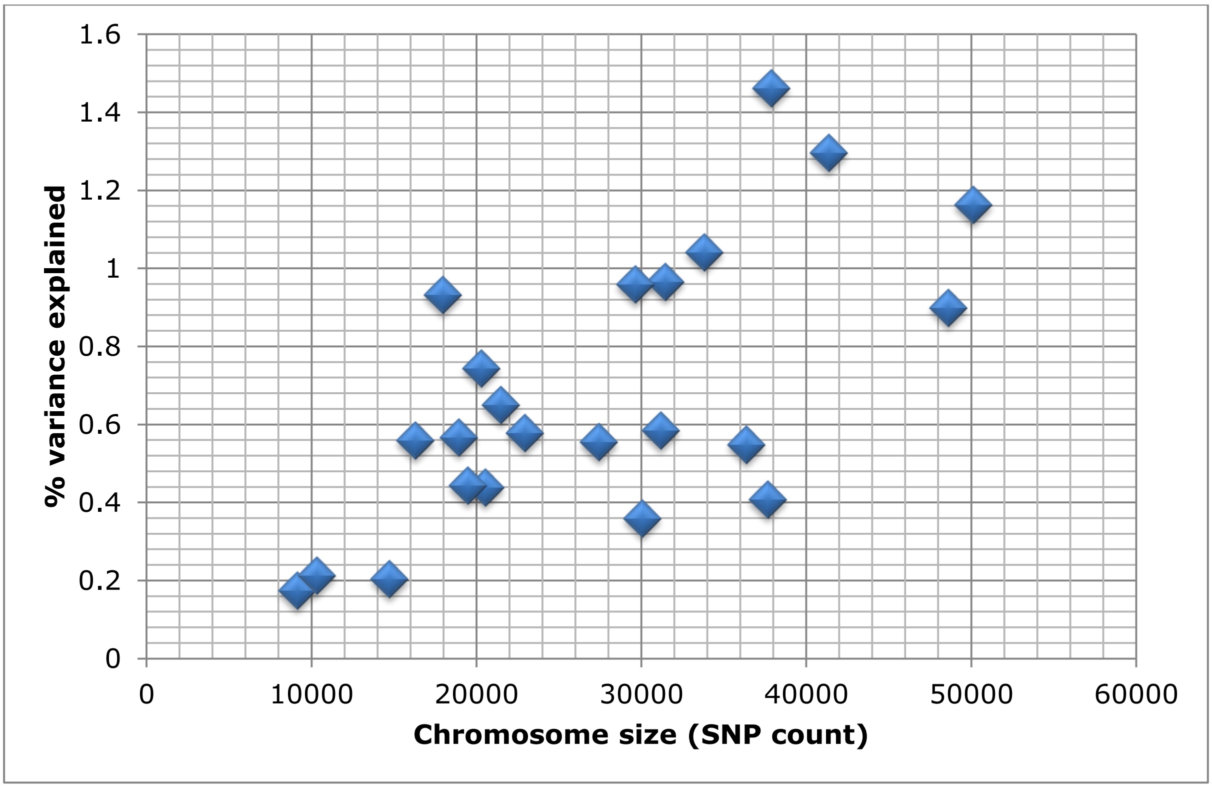

**Figure S2 | Cell and tissue type enrichment estimated using LDSC-SEG**

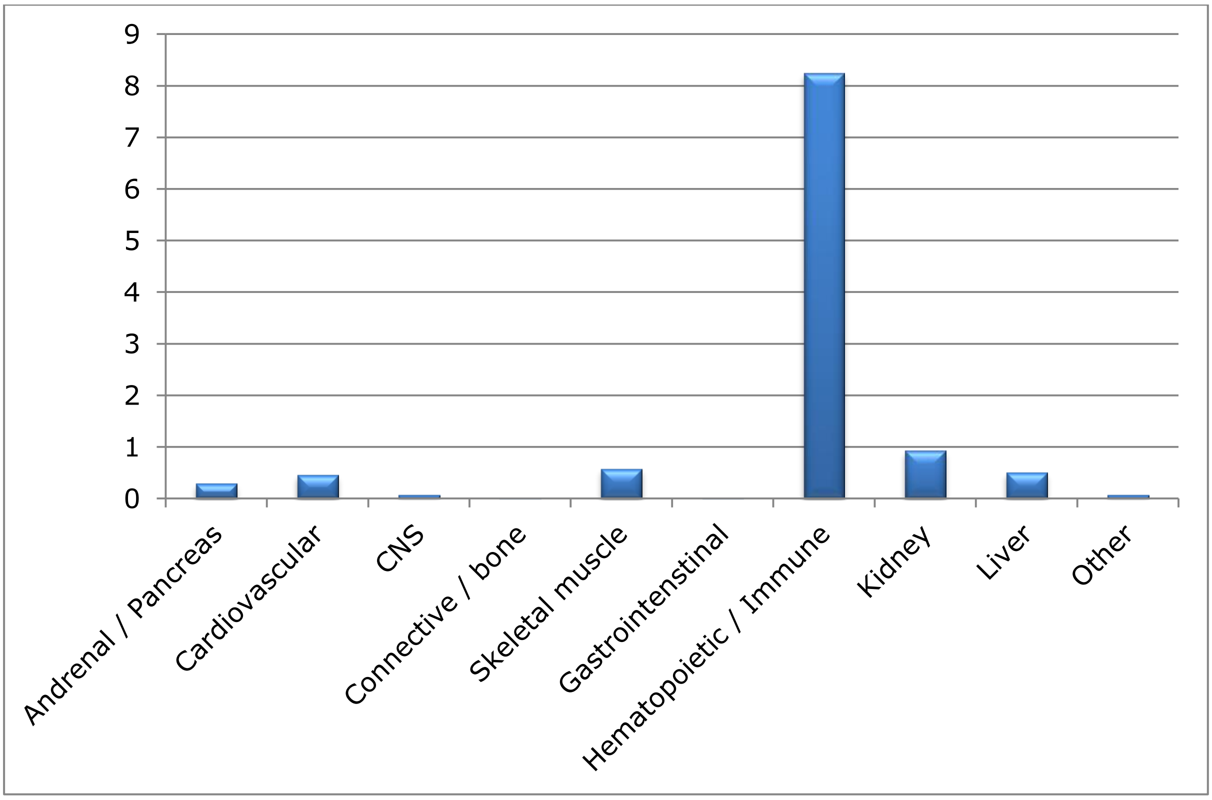

**Supplementary Figure 3.** Differential expression of the *TCL1A* gene in B-lymphocytes with and without the Y chromosome within individual subjects. Error bars indicate the 95% confidence interval of the mean normalized expression of *TCL1A* within each group. To avoid stochastic effects that might occur in estimations using a small number of cells, results are shown for individuals with LOY in at least 10% of the B-lymphocytes and with LOY in more than five individual B-lymphocytes. Within each of the seven individuals (S1-S7) meeting this criteria, *TCL1A* showed a higher expression in the LOY cells compared to normal cells. This suggests that the observed *TCL1A* overexpression in B-lymphocytes without a Y chromosome is independent from the individual genotypes at the lead GWAS-SNP (rs2887399).

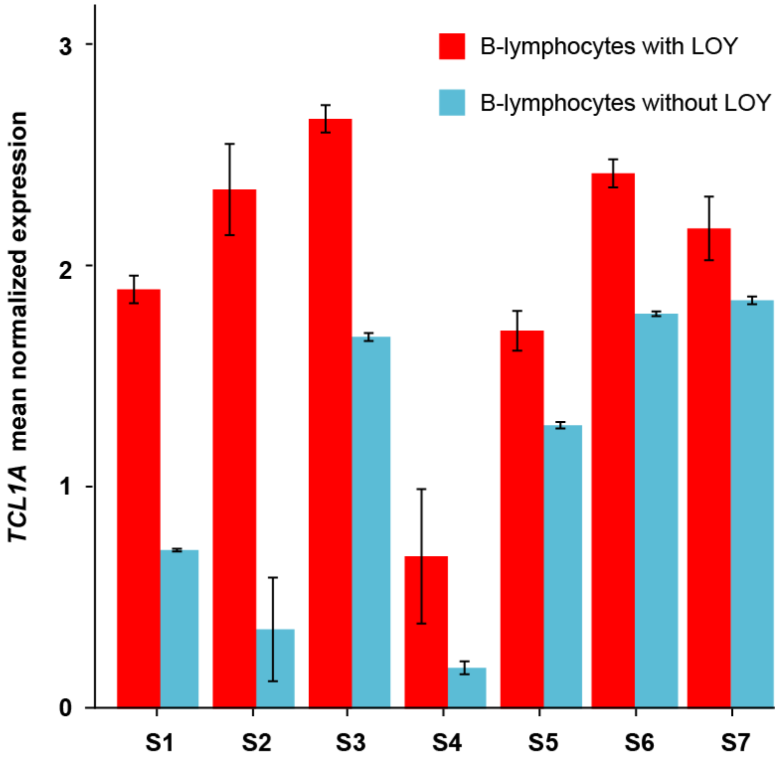

### 1. Consortium Authorship

#### 23andMe Research Team

Michelle Agee, Babak Alipanahi, Adam Auton, Robert K. Bell, Katarzyna Bryc, Sarah L. Elson, Pierre Fontanillas, Nicholas A. Furlotte, David A. Hinds, Karen E. Huber, Aaron Kleinman, Nadia K. Litterman, Matthew H. McIntyre, Joanna L. Mountain, Elizabeth S. Noblin, Carrie A.M. Northover, Steven J. Pitts, J. Fah Sathirapongsasuti, Olga V. Sazonova, Janie F. Shelton, Suyash Shringarpure, Chao Tian, Joyce Y. Tung, Vladimir Vacic, Catherine H. Wilson.

#### The Lung Cancer OncoArray Project (INTEGRAL-ILCCO)

James D McKay[1], Rayjean J Hung[2], David C Christiani[3], Neil E Caporaso[4], Mattias Johansson[1], Stig E Bojesen[5-7], Xifeng Wu[8], Sanjay Shete[8], Loic Le Marchand[9], Demetrios Albanes[4], Heike Bickeböller[10], Melinda C Aldrich[11], Adonina Tardon[12], Gad Rennert[13], M Dawn Teare[14], John K Field[15], Lambertus A Kiemeney[16], Philip Lazarus[17], Stephen Lam[18], Matthew B Schabath[19], Angeline S Andrew[20], Hongbing Shen[21], Yun-Chul Hong[22], Jian-Min Yuan[23], Christopher A Haiman[24], Gary E Goodman[25], H-Erich Wichmann[26–28], Angela Risch[29–33], Susanne M Arnold[34], Shanbeh Zienolddiny[35], Paul Brennan[1], Maria Teresa Landi[4] & Christopher I Amos[36]

[1]International Agency for Research on Cancer, World Health Organization, Lyon, France. [2]Lunenfeld-Tanenbaum Research Institute, Sinai Health System, University of Toronto, Toronto, Ontario, Canada. [3]Department of Environmental Health, Harvard T.H. Chan School of Public Health and Massachusetts General Hospital/ Harvard Medical School, Boston, Massachusetts, USA. [4]Division of Cancer Epidemiology and Genetics, National Cancer Institute, US National Institutes of Health, Bethesda, Maryland, USA. [5]Department of Clinical Biochemistry, Herlev and Gentofte Hospital, Copenhagen University Hospital, Copenhagen, Denmark. [6]Faculty of Health and Medical Sciences, University of Copenhagen, Copenhagen, Denmark. [7]Copenhagen General Population Study, Herlev and Gentofte Hospital, Copenhagen, Denmark. [8]Department of Epidemiology, University of Texas MD Anderson Cancer Center, Houston, Texas, USA. [9]Epidemiology Program, University of Hawaii Cancer Center, Honolulu, Hawaii, USA. [10]Department of Genetic Epidemiology, University Medical Center, Georg August University Göttingen, Göttingen, Germany. [11]Department of Thoracic Surgery, Division of Epidemiology, Vanderbilt University Medical Center, Nashville, Tennessee, USA. [12]University of Oviedo and CIBERESP, Faculty of Medicine, Oviedo, Spain. [13]Clalit National Cancer Control Center at Carmel Medical Center and Technion Faculty of Medicine, Haifa, Israel. [14]School of Health and Related Research, University of Sheffield, Sheffield, UK. [15]Institute of Translational Medicine, University of Liverpool, Liverpool, UK. [16]Departments of Health Evidence and Urology, Radboud University Medical Center, Nijmegen, the Netherlands. [17]Department of Pharmaceutical Sciences, College of Pharmacy, Washington State University, Spokane, Washington, USA. [18]British Columbia Cancer Agency, Vancouver, British Columbia, Canada. [19]Department of Cancer Epidemiology, H. Lee Moffitt Cancer Center and Research Institute, Tampa, Florida, USA. [20]Department of Epidemiology,

Geisel School of Medicine, Hanover, New Hampshire, USA. [21]Department of Epidemiology and Biostatistics, Jiangsu Key Lab of Cancer Biomarkers, Prevention and Treatment, Collaborative Innovation Center for Cancer Personalized Medicine, School of Public Health, Nanjing Medical University, Nanjing, China. [22]Department of Preventive Medicine, Seoul National University College of Medicine, Seoul, Republic of Korea. [23]University of Pittsburgh Cancer Institute, Pittsburgh, Pennsylvania, USA. [24]Department of Preventive Medicine, Keck School of Medicine, University of Southern California Norris Comprehensive Cancer Center, Los Angeles, California, USA. [25]Swedish Medical Group, Seattle, Washington, USA. [26]Institute of Epidemiology II, Helmholtz Zentrum München–German Research Center for Environmental Health, Neuherberg, Germany. [27]Institute of Medical Informatics, Biometry and Epidemiology, Ludwig Maximilians University, Munich, Germany. [28]Institute of Medical Statistics and Epidemiology, Technical University of Munich, Munich, Germany. [29]Research Unit of Molecular Epidemiology, Helmholtz Zentrum München–German Research Center for Environmental Health, Neuherberg, Germany. [30]Thoraxklinik at University Hospital Heidelberg, Heidelberg, Germany. [31]Translational Lung Research Center Heidelberg (TLRC-H), Heidelberg, Germany. [32]German Center for Lung Research (DZL), Heidelberg, Germany. [33]University of Salzburg and Cancer Cluster Salzburg, Salzburg, Austria. [34]Markey Cancer Center, University of Kentucky, Lexington, Kentucky, USA. [35]National Institute of Occupational Health, Oslo, Norway. [36]Biomedical Data Science, Geisel School of Medicine at Dartmouth, Hanover, New Hampshire, USA.

**The PRACTICAL Consortium (http://practical.icr.ac.uk/):**

Rosalind A. Eeles[1,2], Brian E. Henderson[3*], Christopher A. Haiman[3], ZSofia Kote-Jarai[1], Fredrick R. Schumacher[4,5], Ali Amin Al Olama[6,7], Sara Benlloch[6,1], Kenneth Muir[8,9], Sonja I. Berndt[10], David V. Conti[3], Fredrik Wiklund[11], Stephen Chanock[10], Susan Gapstur[12], Victoria L. Stevens[12], Catherine M. Tangen[13], Jyotsna Batra[14,15], Judith Clements[14,15], Australian Prostate Cancer Research Centre BioResource (APCB)[14], Henrik Gronberg[11], Nora Pashayan[16,17], Johanna Schleutker[18,19], Demetrius Albanes[10], Alicja Wolk[20, 21], Catharine West[22], Lorelei Mucci[23], Géraldine Cancel-Tassin[24,25], Stella Koutros[10], Karina Dalsgaard Sorensen[26,27], Eli Marie Grindedal[28], David E. Neal[29,30,31], Freddie C. Hamdy[31], Jenny L. Donovan[32], Ruth C. Travis[33], Robert J. Hamilton[34], Sue Ann Ingles[3], Barry S. Rosenstein[35,36], Yong-Jie Lu[37], Graham G. Giles[38,39], Adam S. Kibel[40], Ana Vega[41], Manolis Kogevinas[42,43,44,45], Kathryn L. Penney[46], Jong Y. Park[47], Janet L. Stanford[48,49], Cezary Cybulski[50], Børge G. Nordestgaard[51,52], Hermann Brenner[53,54,55], Christiane Maier[56], Jeri Kim[57], Esther M. John[58,59], Manuel R. Teixeira[60,61], Susan L. Neuhausen[62], Kim De Ruyck[63], Azad Razack[64], Lisa F. Newcomb[48,65], Davor Lessel[66], Radka Kaneva[67], Nawaid Usmani[68,69], Frank Claessens[70], Paul A. Townsend[71], Manuela Gago-Dominguez[72,73], Monique J. Roobol[74], Florence Menegaux[75], Kay-Tee Khaw[76], Lisa Cannon-Albright[77,78], Hardev Pandha[79], Stephen N. Thibodeau[80]

[*]In memorium

[1] The Institute of Cancer Research, London, UK., [2] Royal Marsden NHS Foundation Trust, London, UK., [3] Department of Preventive Medicine, Keck School of Medicine, University of Southern California/Norris Comprehensive Cancer Center, Los Angeles, CA, USA., [4] Department of Epidemiology and Biostatistics, Case Western Reserve University, Cleveland, OH, USA., [5] Seidman Cancer Center, University Hospitals, Cleveland, OH, USA., [6] Centre for Cancer Genetic Epidemiology, Department of Public Health and Primary Care, University of Cambridge, Strangeways Research Laboratory, Cambridge, UK., [7] University of Cambridge, Department of Clinical Neurosciences, Cambridge, UK., [8] Division of Population Health, Health Services Research and Primary Care, University of Manchester, Manchester, UK., [9] Warwick Medical School, University of Warwick, Coventry, UK., [10] Division of Cancer Epidemiology and Genetics, National Cancer Institute, NIH, Bethesda, MD, USA., [11] Department of Medical Epidemiology and Biostatistics, Karolinska Institute, Stockholm, Sweden., [12] Epidemiology Research Program, American Cancer Society, 250 Williams Street, Atlanta, GA, USA., [13] SWOG Statistical Center, Fred Hutchinson Cancer Research Center, Seattle, WA, USA., [14] Australian Prostate Cancer Research Centre-Qld, Institute of Health and Biomedical Innovation and School of Biomedical Science, Queensland University of Technology, Brisbane, Queensland, Australia., [15] Translational Research Institute, Brisbane, Queensland, Australia., [16] University College London, Department of Applied Health Research, London, UK., [17] Centre for Cancer Genetic Epidemiology, Department of Oncology, University of Cambridge, Strangeways Laboratory, Cambridge, UK., [18] Department of Medical Biochemistry and Genetics, Institute of Biomedicine, University of Turku, Finland., [19] Tyks Microbiology and Genetics, Department of Medical Genetics, Turku University Hospital, Finland., [20] Division of Nutritional Epidemiology, Institute of Environmental Medicine, Karolinska Institutet, Sweden., [21] Department of Surgical Sciences, Uppsala University, Uppsala, Sweden., [22] Division of Cancer Sciences, University of Manchester, Manchester Academic Health Science Centre, Radiotherapy Related Research, Manchester NIHR Biomedical Research Centre, The Christie Hospital NHS Foundation Trust, Manchester, UK., [23] Department of Epidemiology, Harvard T.H Chan School of Public Health, Boston, MA, USA., [24] CeRePP, Tenon Hospital, Paris, France., [25] UPMC Sorbonne Universites, GRC N°5 ONCOTYPE-URO, Tenon Hospital, Paris, France., [26] Department of Molecular Medicine, Aarhus University Hospital, Denmark., [27] Department of Clinical Medicine, Aarhus University, Denmark., [28] Department of Medical Genetics, Oslo University Hospital, Norway., [29] University of Cambridge, Department of Oncology, Addenbrooke's Hospital, Cambridge, UK., [30] Cancer Research UK Cambridge Research Institute, Li Ka Shing Centre, Cambridge, UK., [31] Nuffield Department of Surgical Sciences, University of Oxford, Oxford, UK, Faculty of Medical Science, University of Oxford, John Radcliffe Hospital, Oxford, UK., [32] School of Social and Community Medicine, University of Bristol, Bristol, UK., [33] Cancer Epidemiology Unit, Nuffield Department of Population Health University of Oxford, Oxford, UK., [34] Dept. of Surgical Oncology, Princess Margaret Cancer Centre, Toronto, Canada., [35] Department of Radiation Oncology, Icahn School of Medicine at Mount Sinai, New York, NY, USA., [36] Department of Genetics and Genomic Sciences, Icahn School of Medicine at Mount Sinai,

New York, NY, USA., [37] Centre for Molecular Oncology, Barts Cancer Institute, Queen Mary University of London, John Vane Science Centre, London, UK., [38] Cancer Epidemiology & Intelligence Division, The Cancer Council Victoria, Melbourne, Victoria, Australia., [39] Centre for Epidemiology and Biostatistics, Melbourne School of Population and Global Health, The University of Melbourne, Melbourne, Australia., [40] Division of Urologic Surgery, Brigham and Womens Hospital, Boston, MA, USA., [41] Fundación Pública Galega de Medicina Xenómica-SERGAS, Grupo de Medicina Xenómica, CIBERER, IDIS, Santiago de Compostela, Spain., [42] Centre for Research Environmental Epidemiology (CREAL), Barcelona Institute for Global Health (ISGlobal), Barcelona, Spain., [43] CIBER Epidemiología y Salud Pública (CIBERESP), Madrid, Spain., [44] IMIM (Hospital del Mar Research Institute), Barcelona, Spain., [45] Universitat Pompeu Fabra (UPF), Barcelona, Spain., [46] Channing Division of Network Medicine, Department of Medicine, Brigham and Women's Hospital/Harvard Medical School, Boston, MA, USA., [47] Department of Cancer Epidemiology, Moffitt Cancer Center, Tampa, USA., [48] Division of Public Health Sciences, Fred Hutchinson Cancer Research Center, Seattle, Washington, USA., [49] Department of Epidemiology, School of Public Health, University of Washington, Seattle, Washington, USA., [50] International Hereditary Cancer Center, Department of Genetics and Pathology, Pomeranian Medical University, Szczecin, Poland., [51] Faculty of Health and Medical Sciences, University of Copenhagen, Denmark., [52] Department of Clinical Biochemistry, Herlev and Gentofte Hospital, Copenhagen University Hospital, Herlev, Denmark., [53] Division of Clinical Epidemiology and Aging Research, German Cancer Research Center (DKFZ), Heidelberg, Germany., [54] German Cancer Consortium (DKTK), German Cancer Research Center (DKFZ), Heidelberg, Germany., [55] Division of Preventive Oncology, German Cancer Research Center (DKFZ) and National Center for Tumor Diseases (NCT), Heidelberg, Germany., [56] Institute for Human Genetics, University Hospital Ulm, Ulm, Germany., [57] The University of Texas MD Anderson Cancer Center, Department of Genitourinary Medical Oncology, Houston, TX, USA., [58] Cancer Prevention Institute of California, Fremont, CA, USA., [59] Department of Health Research & Policy (Epidemiology) and Stanford Cancer Institute, Stanford University School of Medicine, Stanford, CA , USA., [60] Department of Genetics, Portuguese Oncology Institute of Porto, Porto, Portugal., [61] Biomedical Sciences Institute (ICBAS), University of Porto, Porto, Portugal., [62] Department of Population Sciences, Beckman Research Institute of the City of Hope, Duarte, CA, USA., [63] Ghent University, Faculty of Medicine and Health Sciences, Basic Medical Sciences, Gent, Belgium., [64] Department of Surgery, Faculty of Medicine, University of Malaya, Kuala Lumpur, Malaysia., [65] Department of Urology, University of Washington, Seattle, WA, USA., [66] Institute of Human Genetics, University Medical Center Hamburg-Eppendorf, Hamburg, Germany., [67] Molecular Medicine Center, Department of Medical Chemistry and Biochemistry, Medical University, Sofia, Bulgaria., [68] Department of Oncology, Cross Cancer Institute, University of Alberta, Edmonton, Alberta, Canada., [69] Division of Radiation Oncology, Cross Cancer Institute, Edmonton, Alberta, Canada., [70] Molecular Endocrinology Laboratory, Department of Cellular and Molecular Medicine, KU Leuven, Leuven, Belgium., [71] Institute of Cancer Sciences, Manchester Cancer Research

Centre, University of Manchester, Manchester Academic Health Science Centre, St Mary's Hospital, Manchester, UK., [72] Genomic Medicine Group, Galician Foundation of Genomic Medicine, Instituto de Investigacion Sanitaria de Santiago de Compostela (IDIS), Complejo Hospitalario Universitario de Santiago, Servicio Galego de Saúde, SERGAS, Santiago De Compostela, Spain., [73] University of California San Diego, Moores Cancer Center, La Jolla, CA, USA., [74] Department of Urology, Erasmus University Medical Center, Rotterdam, the Netherlands., [75] Cancer & Environment Group, Center for Research Epidemiology and Population Health (CESP), INSERM, University Paris-Sud, University Paris-Saclay, Villejuif, France., [76] Clinical Gerontology Unit, University of Cambridge, Cambridge, UK., [77] Division of Genetic Epidemiology, Department of Medicine, University of Utah School of Medicine, Salt Lake City, Utah, USA., [78] George E. Wahlen Department of Veterans Affairs Medical Center, Salt Lake City, UT, USA., [79] The University of Surrey, Guildford, Surrey, UK., [80] Department of Laboratory Medicine and Pathology, Mayo Clinic, Rochester, MN, USA.

**Consortium of Investigators of Modifiers of BRCA1/2 (CIMBA)**

Irene L. Andrulis[1,2], Antonis C. Antoniou[3], Banu K. Arun[4], Javier Benitez[5,6], Ake Borg[7], Trinidad Caldés[8], Maria A. Caligo[9], Georgia Chenevix-Trench[10], Wendy K. Chung[11], Kathleen B.M. Claes[12], Fergus J. Couch[13], Mary B. Daly[14], Peter Devilee[15,16], Orland Diez[17], Douglas F. Easton[3,18], Eitan Friedman[19,20], Patricia A. Ganz[21], Judy Garber[22], Andrew K. Godwin[23], David E. Goldgar[24], Mark H. Greene[25], Ute Hamann[26], Frans B.L. Hogervorst[27], John L. Hopper[28], Peter J. Hulick[29,30], Evgeny N. Imyanitov[31], Claudine Isaacs[32], Anna Jakubowska[33,3,4], Paul James[35,36], Ramunas Janavicius[37], Esther M. John[38-40], Beth Y. Karlan[41], Conxi Lazaro[42], Fabienne Lesueur[43-46], Austin Miller[47], Marco Montagna[48], Katherine L. Nathanson[49], Susan L. Neuhausen[50], Heli Nevanlinna[51], Finn C. Nielsen[52], Liene Nikitina-Zake[53], Robert L. Nussbaum[54], Kenneth Offit[55,56], Edith Olah[57], Olufunmilayo I. Olopade[58], Paolo Radice[59], Johanna Rantala[60], Gadi Rennert[61], Harvey A. Risch[62], Kristin Sanden[63], Rita K. Schmutzler[64,65], Jacques Simard[66], Christian F. Singer[67], Penny Soucy[66], Melissa C. Southey[68,69], Manuel R. Teixeira[70,71], Mary Beth Terry[72], Mads Thomassen[73], Darcy L. Thull[74], Marc Tischkowitz[75,76], Amanda E. Toland[77], Nadine Tung[78], Elizabeth J. van Rensburg[79], Ana Vega[6,80,81], Barbara Wappenschmidt[64,65], Jeffrey N. Weitzel[82], Drakoulis Yannoukakos[83]

Fred A. Litwin Center for Cancer Genetics, Lunenfeld-Tanenbaum Research Institute of Mount Sinai Hospital, Toronto, ON, Canada, 2. Department of Molecular Genetics, University of Toronto, Toronto, ON, Canada, 3. Centre for Cancer Genetic Epidemiology, Department of Public Health and Primary Care, University of Cambridge, UK, 4. Department of Breast Medical Oncology, University of Texas MD Anderson Cancer Center, Houston, TX, USA, 5. Human Cancer Genetics Programme, Spanish National Cancer Research Centre (CNIO), Madrid, Spain., 6. Biomedical Network on Rare Diseases (CIBERER), Madrid, Spain., 7. Department of Oncology, Lund University and Skåne University Hospital, Lund, Sweden.,

8. Medical Oncology Department, Hospital Clínico San Carlos, Instituto de Investigación Sanitaria San Carlos (IdISSC), Centro Investigación Biomédica en Red de Cáncer (CIBERONC), Madrid, Spain., 9. Section of Molecular Genetics, Dept. of Laboratory Medicine, University Hospital of Pisa, Italy., 10. Department of Genetics and Computational Biology, QIMR Berghofer Medical Research Institute, Brisbane, Queensland, Australia., 11. Departments of Pediatrics and Medicine, Columbia University, New York, NY, USA., 12. Centre for Medical Genetics, Ghent University, Ghent, Belgium., 13. Department of Laboratory Medicine and Pathology, Mayo Clinic, Rochester, MN, USA., 14. Department of Clinical Genetics, Fox Chase Cancer Center, Philadelphia, PA, USA., 15. Department of Pathology, Leiden University Medical Center, Leiden, The Netherlands., 16. Department of Human Genetics, Leiden University Medical Center, Leiden, The Netherlands., 17. Oncogenetics Group, Clinical and Molecular Genetics Area, Vall d'Hebron Institute of Oncology (VHIO), University Hospital, Vall d'Hebron, Barcelona, Spain., 18. Centre for Cancer Genetic Epidemiology, Department of Oncology, University of Cambridge, UK., 19. The Susanne Levy Gertner Oncogenetics Unit, Chaim Sheba Medical Center, Ramat Gan, Israel., 20. Sackler Faculty of Medicine, Tel Aviv University, Ramat Aviv, Israel, 21. Schools of Medicine and Public Health, Division of Cancer Prevention & Control Research, Jonsson Comprehensive Cancer Centre, UCLA, Los Angeles, CA, USA., 22. Cancer Risk and Prevention Clinic, Dana-Farber Cancer Institute, Boston, MA, USA., 23. Department of Pathology and Laboratory Medicine, Kansas University Medical Center, Kansas City, KS, USA., 24. Department of Dermatology, Huntsman Cancer Institute, University of Utah School of Medicine, Salt Lake City, UT, USA., 25. Clinical Genetics Branch, Division of Cancer Epidemiology and Genetics, National Cancer Institute, Bethesda, MD, USA., 26. Molecular Genetics of Breast Cancer, German Cancer Research Center (DKFZ), Heidelberg, Germany., 27. Family Cancer Clinic, The Netherlands Cancer Institute - Antoni van Leeuwenhoek hospital, Amsterdam, The Netherlands., 28. Centre for Epidemiology and Biostatistics, Melbourne School of Population and Global Health, The University of Melbourne, Melbourne, Victoria, Australia., 29. Center for Medical Genetics, NorthShore University HealthSystem, Evanston, IL, USA., 30. The University of Chicago Pritzker School of Medicine, Chicago, IL, USA., 31. N.N. Petrov Institute of Oncology, St. Petersburg, Russia., 32. Lombardi Comprehensive Cancer Center, Georgetown University, Washington, DC, USA., 33. Department of Genetics and Pathology, Pomeranian Medical University, Szczecin, Poland., 34. Independent Laboratory of Molecular Biology and Genetic Diagnostics, Pomeranian Medical University, Szczecin, Poland., 35. Parkville Familial Cancer Centre, Peter MacCallum Cancer Center, Melbourne, Victoria Australia., 36. Sir Peter MacCallum Department of Oncology, The University of Melbourne, Melbourne, Victoria Australia., 37. Hematology, oncology and transfusion medicine center, Dept. of Molecular and Regenerative Medicine, Vilnius University Hospital Santariskiu Clinics, Vilnius, Lithuania., 38. Department of Epidemiology, Cancer Prevention Institute of California, Fremont, CA, USA., 39. Department of Health Research and Policy - Epidemiology. Stanford University School of Medicine, Stanford, CA, USA., 40. Department of Biomedical Data Science, Stanford University School of Medicine, Stanford, CA, USA)., 41. Women's Cancer Program at the

Samuel Oschin Comprehensive Cancer Institute, Cedars-Sinai Medical Center, Los Angeles, CA, USA., 42. Molecular Diagnostic Unit, Hereditary Cancer Program, IDIBELL (Bellvitge Biomedical Research Institute), Catalan Institute of Oncology, CIBERONC, Barcelona, Spain., 43. Genetic Epidemiology of Cancer team, Inserm U900, Paris France., 44. Service de Génétique, Institut Curie, Paris, France., 45. PSL University, Paris, France., 46. Mines ParisTech, Fontainebleau, France., 47. NRG Oncology, Statistics and Data Management Center, Roswell Park Cancer Institute, Buffalo, NY, USA., 48. Immunology and Molecular Oncology Unit, Veneto Institute of Oncology IOV - IRCCS, Padua, Italy., 49. Department of Medicine, Abramson Cancer Center, Perelman School of Medicine at the University of Pennsylvania, Philadelphia, PA, USA., 50. Department of Population Sciences, Beckman Research Institute of City of Hope, Duarte, CA, USA., 51. Department of Obstetrics and Gynecology, Helsinki University Hospital, University of Helsinki, Finland., 52. Center for Genomic Medicine, Rigshospitalet, Copenhagen University Hospital, Copenhagen, Denmark., 53. Latvian Biomedical Research and Study Centre, Riga, Latvia., 54. Cancer Genetics and Prevention Program, University of California San Francisco, San Francisco, CA, USA., 55. Clinical Genetics Research Lab, Department of Cancer Biology and Genetics, Memorial Sloan-Kettering Cancer Center, New York, NY, USA., 56. Clinical Genetics Service, Department of Medicine, Memorial Sloan-Kettering Cancer Center, New York, NY, USA., 57. Department of Molecular Genetics, National Institute of Oncology, Budapest, Hungary., 58. Center for Clinical Cancer Genetics, The University of Chicago, Chicago, IL, USA., 59. Unit of Molecular Bases of Genetic Risk and Genetic Testing, Department of Research, Fondazione IRCCS (Istituto Di Ricovero e Cura a Carattere Scientifico) Istituto Nazionale dei Tumori (INT), Milan, Italy., 60. Clinical Genetics, Karolinska Institutet, Stockholm, Sweden., 61. Clalit National Cancer Control Center, Carmel Medical Center and Technion Faculty of Medicine, Haifa, Israel., 62. Chronic Disease Epidemiology, Yale School of Public Health, New Haven, CT, USA., 63. City of Hope Clinical Cancer Genetics Community Research Network, Duarte, CA, USA., 64. Center for Hereditary Breast and Ovarian Cancer, University Hospital of Cologne, Cologne, Germany., 65. Center for Molecular Medicine Cologne (CMMC), University of Cologne, Cologne, Germany., 66. Genomics Center, Centre Hospitalier Universitaire de Québec Research Center, Laval University, Québec City, QC, Canada., 67. Dept of OB/GYN and Comprehensive Cancer Center, Medical University of Vienna, Vienna, Austria., 68. Precision Medicine, School of Clinical Sciences at Monash Health, Monash University, Clayton, Victoria, Australia., 69. Department of Clinical Pathology, The University of Melbourne, Melbourne, Victoria, Australia., 70. Department of Genetics, Portuguese Oncology Institute, Porto, Portugal., 71. Biomedical Sciences Institute (ICBAS), University of Porto, Porto, Portugal., 72. Department of Epidemiology, Mailman School of Public Health, Columbia University, New York, NY, USA., 73. Department of Clinical Genetics, Odense University Hospital, Odence, Denmark, 74. Department of Medicine, Magee-Womens Hospital, University of Pittsburgh School of Medicine, Pittsburgh, PA, USA., 75. Program Cancer Genetics, Departments of Human Genetics and Oncolog, McGill University, Montréal, QC, Canada., 76. Department of Medical Genetics, University of Cambridge,

Addenbrooke's Hosptital, Cambridge, UK., 77. Department of Cancer Biology and Genetics, The Ohio State University, Columbus, OH, USA., 78. Department of Medical Oncology, Beth Israel Deaconess Medical Center, Boston, MA, USA., 79. Department of Genetics, University of Pretoria, Arcadia, South Africa., 80. Fundación Pública Galega Medicina Xenómica, Santiago De Compostela, Spain., 81. Instituto de Investigación Sanitaria de Santiago de Compostela, Santiago De Compostela, Spain., 82. Clinical Cancer Genetics, City of Hope, Duarte, CA, USA., 83. Molecular Diagnostics Laboratory, INRASTES, National Centre for Scientific Research 'Demokritos', Athens, Greece.

**Kidney Cancer GWAS Meta-Analysis Project**

Jonathan N. Hofmann[1], Robert Carreras-Torres[2], Kevin M. Brown[1], Mattias Johansson[2], Zhaoming Wang[3], Matthieu Foll[2], Peng Li[2], Nathaniel Rothman[1], Sharon A. Savage[1], Valerie Gaborieau[2], James D. McKay[2], Yuanqing Ye[4], Marc Henrion[5], Fiona Bruinsma[6], Susan Jordan[7,8], Gianluca Severi[6,9,10,11], Kristian Hveem[12], Lars J. Vatten[13], Tony Fletcher[14], Kvetoslava Koppova[15], Susanna C. Larsson[16], Alicja Wolk[16], Rosamonde E. Banks[17], Peter J. Selby[17], Gabriella Andreotti[1], Laura E. Beane Freeman[1], Stella Koutros[1], Demetrius Albanes[1], Satu Mannisto[18], Stephanie Weinstein[1], Peter E. Clark[19], Todd E. Edwards[19], Loren Lipworth[19], Susan M. Gapstur[20], Victoria L. Stevens[20], Hallie Carol[21], Matthew L. Freedman[21], Mark M. Pomerantz[21], Eunyoung Cho[22], Peter Kraft[23], Mark A. Preston[24], Kathryn M. Wilson[23], J. Michael Gaziano[24,25], Howard S. Sesso[23], Amanda Black[1], Neal D. Freedman[1], Wen-Yi Huang[1], John G. Anema[26], Richard J. Kahnoski[26], Brian R. Lane[26,27], Sabrina L. Noyes[28], David Petillo[28], Leandro M. Colli[1], Joshua N. Sampson[1], Celine Besse[29], Helene Blanche[30], Anne Boland[29], Laurie Burdette[1], Egor Prokhortchouk[31,32], Konstantin G. Skryabin[31,32], Meredith Yeager[1], Mirjana Mijuskovic[33], Miodrag Ognjanovic[34], Lenka Foretova[35], Ivana Holcatova[36], Vladimir Janout[37], Dana Mates[38], Anush Mukeriya[39], Stefan Rascu[40], David Zaridze[39], Vladimir Bencko[41], Cezary Cybulski[42], Eleonora Fabianova[15], Viorel Jinga[40], Jolanta Lissowska[43], Jan Lubinski[42], Marie Navratilova[35], Peter Rudnai[44], Neonila Szeszenia-Dabrowska[45], Simone Benhamou[46,47], Geraldine Cancel-Tassin[48,49], Olivier Cussenot[48,49,50], H.B(as). Bueno-de-Mesquita[51,52,53,54], Federico Canzian[55], Eric J. Duell[56], Börje Ljungberg[57], Raviprakash T. Sitaram[57], Ulrike Peters[58], Emily White[58], Garnet L. Anderson[58], Lisa Johnson[58], Juhua Luo[59], Julie Buring[23], I-Min Lee[23,24], Wong-Ho Chow[4], Lee E. Moore[1], Christopher Wood[60], Timothy Eisen[61], James Larkin[62], Toni K. Choueiri[21], G. Mark Lathrop[63], Bin Tean Teh[28], Jean-Francois Deleuze[29,30], Xifeng Wu[4], Richard S. Houlston[64], Paul Brennan[2], Ghislaine Scelo[2], Mark P. Purdue[1]

[1]Division of Cancer Epidemiology and Genetics, National Cancer Institute, National Institutes of Health, Department Health and Human Services, Bethesda, MS, USA, [2]International Agency for Research on Cancer (IARC), Lyon, France, [3]St. Jude Children's Research Hospital, Memphis, TN, USA, [4]Department of Epidemiology, Division of Cancer Prevention and Population Sciences, The University of Texas MD Anderson Cancer Center, Houston, TX, USA, [5]Icahn School of Medicine, New York, NY, USA, [6]Cancer Epidemiology and Intelligence Division, Cancer Council Victoria, Melbourne, Australia, [7]QIMR Berghofer Medical Research Institute, Herston, Queensland, Australia, [8]School of Public Health, The University of

Queensland, Brisbane, Australia, [9]Centre for Epidemiology and Biostatistics, Melbourne School of Population and Global Health, The University of Melbourne, Carlton, Australia, [10]Human Genetics Foundation (HuGeF), Torino, Italy, [11]Centre de Recherche en Épidémiologie et Santé des Populations, Université Paris-Saclay, UPS, USQ, Gustave Roussy, Villejuif, France, [12]HUNT Research Centre, Department of Public Health and General Practice, Norwegian University of Science and Technology, Levanger, Sweden, [13]Department of Public Health and General Practice, Faculty of Medicine, Norwegian University of Science and Technology, Trondheim, Norway, [14]London School of Hygiene and Tropical Medicine, University of London, London, UK, [15]Regional Authority of Public Health Banska Bystrica, Banska Bystrica, Slovakia, [16]Institute of Environmental Medicine, Karolinska Institutet, Stockholm, Sweden, [17]Leeds Institute of Cancer and Pathology, University of Leeds, Cancer Research Building, St James' University Hospital, Leeds, UK, [18]National Institute for Health and Welfare, Helsinki, Finland, [19]Vanderbilt-Ingram Cancer Center, Nashville, TN, USA, [20]American Cancer Society, Atlanta, GA, USA, [21]Dana-Farber Cancer Institute, Boston, MA, USA, [22]Brown University, Providence, RI, USA, [23]Harvard T.H. Chan School of Public Health, Boston, MA, USA, [24]Brigham and Women's Hospital, Boston, MA, USA, [25]Veterans Administration, Boston, MA, USA, [26]Division of Urology, Spectrum Health, Grand Rapids, MI, USA, [27]College of Human Medicine, Michigan State University, Grand Rapids, MI, USA, [28]Van Andel Research Institute, Center for Cancer Genomics and Quantitative Biology, Grand Rapids, MI, USA, [29]Centre National de Recherche en Genomique Humaine (CNRGH), Institut de biologie François Jacob, Commissariat à l'Energie Atomique et aux Energies Alternatives, Evry, France, [30]Fondation Jean Dausset-Centre d'Etude du Polymorphisme Humain, Paris, France, [31]Center 'Bioengineering' of the Russian Academy of Sciences, Moscow, Russian Federation, [32]Kurchatov Scientific Center, Moscow, Russian Federation, [33]Clinic for Nephrology, Military Medical Academy, Belgrade, Serbia, [34]International Organization for Cancer Prevention and Research (IOCPR), Belgrade, Serbia, [35]Department of Cancer Epidemiology and Genetics, Masaryk Memorial Cancer Institute, Brno, Czech Republic, [36]Second Faculty of Medicine, Institute of Public Health and Preventive Medicine, Charles University, Prague, Czech Republic, [37]Department of Preventive Medicine, Faculty of Medicine, Palacky University, Czech Republic, [38]National Institute of Public Health, Bucharest, Romania, [39]Russian N.N. Blokhin Cancer Research Centre, Moscow, Russian Federation, [40]Carol Davila University of Medicine and Pharmacy, Th. Burghele Hospital, Bucharest, Romania, [41]First Faculty of Medicine, Institute of Hygiene and Epidemiology, Charles University, Prague, Czech Republic, [42]International Hereditary Cancer Center, Department of Genetics and Pathology, Pomeranian Medical University, Szczecin, Poland, [43]The M Sklodowska-Curie Cancer Center and Institute of Oncology, Warsaw, Poland, [44]National Public Health Center, National Directorate of Environmental Health, Budapest, Hungary, [45]Department of Epidemiology, Institute of Occupational Medicine, Lodz, Poland, [46]INSERM U946, Paris, France, [47]CNRS UMR8200, Institute Gustave Roussy, Villejuif, France, [48]CeRePP, Paris, France, [49]UPMC Univ Paris 06, Institut Universitaire de Cancérologie, Paris, France, [50]AP-HP, Department of Urology, Hopitaux Universitaires Est Parisien Tenon, Paris, France, [51]Department for Determinants of Chronic Diseases (DCD), National Institute for Public Health and the Environment (RIVM), Bilthoven, The Netherlands, [52]Department of Gastroenterology and Hepatology, University Medical Centre, Utrecht, The Netherlands, [53]Department of Epidemiology and Biostatistics, The School of Public Health, Imperial College London, St Mary's Campus, Norfolk Place, London, UK, [54]Department of Social & Preventive Medicine, Faculty of Medicine, University of Malaya, Pantai Valley, Kuala

Lumpur, Malaysia, [55]Genomic Epidemiology Group, German Cancer Research Center (DKFZ), Heidelberg, Germany, [56]Unit of Nutrition and Cancer, Cancer Epidemiology Research Program, Catalan Institute of Oncology (ICO-IDIBELL), Barcelona, Spain, [57]Department of Surgical and Perioperative Sciences, Urology and Andrology, Umeå University, Umeå, Sweden, [58]Fred Hutchinson Cancer Research Center, Seattle, WA, USA, [59]Department of Epidemiology and Biostatistics, School of Public Health Indiana University Bloomington, Bloomington, IN, USA, [60]Department of Urology, The University of Texas MD Anderson Cancer Center, Houston, TX, USA, [61]University of Cambridge, Cambridge, UK, [62]Royal Marsden NHS Foundation Trust, London, UK, [63]McGill University and Genome Quebec Innovation Centre, Montreal, QC, Canada, [64]The Institute of Cancer Research, London, UK

**The Endometrial Cancer Association Consortium**

Tracy A. O'Mara[1], Amanda B. Spurdle[1], Deborah J. Thompson[2], Frederic Amant[3], Daniela Annibali[3], Katie Ashton[4-6], John Attia[4, 7], Paul L. Auer[8, 9], Matthias W. Beckmann[10], Amanda Black[11], Louise Brinton[11], Daniel D. Buchanan[12-15], Stephen J. Chanock[16], Chu Chen[17], Maxine M. Chen[18], Timothy H.T. Cheng[19], Linda S. Cook[20, 21], Marta Crous-Bous[18, 22], Kamila Czene[23], Immaculata De Vivo[18, 22], Jeroen Depreeuw[3, 24, 25], Jennifer Anne Doherty[26], Thilo Dörk[27], Sean C. Dowdy[28], Alison M. Dunning[29], Matthias Dürst[30], Douglas F. Easton[2, 29], Arif B. Ekici[31], Peter A. Fasching[10, 32], Brooke L. Fridley[33], Christine M. Friedenreich[21], Montserrat García-Closas[16, 34], Mia M. Gaudet[35], Graham G. Giles[13, 36, 37], Dylan M. Glubb[1], Ellen L. Goode[38], Maggie Gorman[19], Christopher A. Haiman[39], Per Hall[23, 40], Susan E. Hankinson[22, 41], Patricia Hartge[11], Catherine S. Healey[29], Alexander Hein[10], Peter Hillemanns[27], Shirley Hodgson[42], Erling Hoivik[43, 44], Elizabeth G. Holliday[4, 7], David J. Hunter[18, 45, 46], Angela Jones[19], Peter Kraft[18, 45], Camilla Krakstad[43, 44], Diether Lambrechts[25, 47], Loic Le Marchand[48], Xiaolin Liang[49], Annika Lindblom[50, 51], Jolanta Lissowska[52], Jirong Long[53], Lingeng Lu[54], Anthony M. Magliocco[55], Lynn Martin[56], Mark McEvoy[7], Roger L. Milne[13, 36, 57], Miriam Mints[58], Rami Nassir[59], Irene Orlow[49], Geoffrey Otton[60], Claire Palles[19], Paul D.P. Pharoah[2, 29], Loreall Pooler[39], Jennifer Prescott[22], Tony Proietto[60], Timothy R. Rebbeck[61, 62], Stefan P. Renner[63], Harvey A. Risch[54], Matthias Rübner[63], Ingo Runnebaum[30], Carlotta Sacerdote[64, 65], Gloria E. Sarto[66], Fredrick Schumacher[67], Rodney J. Scott[4, 6, 68], V. Wendy Setiawan[39], Mitul Shah[29], Xin Sheng[39], Xiao-Ou Shu[53], Melissa C. Southey[12, 57], Emma Tham[50, 69], Ian Tomlinson[19, 56], Jone Trovik[43, 44], Constance Turman[18], David Van Den Berg[39], Adriaan Vanderstichele[70], Zhaoming Wang[11], Penelope M. Webb[71], Nicolas Wentzensen[11], Henrica M.J. Werner[43, 44], Stacey J. Winham[72], Lucy Xia[39], Yong-Bing Xiang[73], Hannah P. Yang[11], Herbert Yu[48], Wei Zheng[53]

[1] Department of Genetics and Computational Biology, QIMR Berghofer Medical Research Institute, Brisbane, Queensland, Australia. [2] Centre for Cancer Genetic Epidemiology, Department of Public Health and Primary Care, University of Cambridge, Cambridge, UK. [3] Department of Obstetrics and Gynecology, Division of Gynecologic Oncology, University Hospitals KU Leuven, University of Leuven, Leuven, Belgium. [4] Hunter Medical Research Institute, John Hunter Hospital, Newcastle, New South Wales, Australia. [5] Centre for Information Based Medicine, University of Newcastle, Callaghan, New South Wales, Australia. [6] Discipline of Medical Genetics, School of Biomedical Sciences and Pharmacy,

Faculty of Health, University of Newcastle, Callaghan, New South Wales, Australia. [7] Centre for Clinical Epidemiology and Biostatistics, School of Medicine and Public Health, University of Newcastle, Callaghan, New South Wales, Australia. [8] Cancer Prevention Program, Fred Hutchinson Cancer Research Center, Seattle, WA, USA. [9] Zilber School of Public Health, University of Wisconsin-Milwaukee, Milwaukee, WI, USA. [10] Department of Gynecology and Obstetrics, Comprehensive Cancer Center ER-EMN, University Hospital Erlangen, Friedrich-Alexander-University Erlangen-Nuremberg, Erlangen, Germany. [11] Division of Cancer Epidemiology and Genetics, National Cancer Institute, Bethesda, MD, USA. [12] Department of Clinical Pathology, The University of Melbourne, Melbourne, Victoria, Australia. [13] Centre for Epidemiology and Biostatistics, Melbourne School of Population and Global Health, The University of Melbourne, Melbourne, Victoria, Australia. [14] Genetic Medicine and Family Cancer Clinic, Royal Melbourne Hospital, Parkville, Victoria, Australia. [15] University of Melbourne Centre for Cancer Research, Victorian Comprehensive Cancer Centre, Parkville, Victoria, Australia. [16] Division of Cancer Epidemiology and Genetics, National Cancer Institute, National Institutes of Health, Department of Health and Human Services, Bethesda, MD, USA. [17] Epidemiology Program, Fred Hutchinson Cancer Research Center, Seattle, WA, USA. [18] Department of Epidemiology, Harvard T.H. Chan School of Public Health, Boston, MA, USA. [19] Wellcome Trust Centre for Human Genetics and Oxford NIHR Biomedical Research Centre, University of Oxford, Oxford, UK. [20] University of New Mexico Health Sciences Center, University of New Mexico, Albuquerque, NM, USA. [21] Department of Cancer Epidemiology and Prevention Research, Alberta Health Services, Calgary, AB, Canada. [22] Channing Division of Network Medicine, Department of Medicine, Brigham and Women's Hospital, Harvard Medical School, Boston, MA, USA. [23] Department of Medical Epidemiology and Biostatistics, Karolinska Institutet, Stockholm, Sweden. [24] Vesalius Research Center, VIB, Leuven, Belgium. [25] Laboratory for Translational Genetics, Department of Human Genetics, University of Leuven, Leuven, Belgium. [26] Huntsman Cancer Institute, Department of Population Health Sciences, University of Utah, Salt Lake City, UT, USA. [27] Gynaecology Research Unit, Hannover Medical School, Hannover, Germany. [28] Department of Obstetrics and Gynecology, Division of Gynecologic Oncology, Mayo Clinic, Rochester, MN, USA. [29] Centre for Cancer Genetic Epidemiology, Department of Oncology, University of Cambridge, Cambridge, UK. [30] Department of Gynaecology, Jena University Hospital - Friedrich Schiller University, Jena, Germany. [31] Institute of Human Genetics, University Hospital Erlangen, Friedrich-Alexander University Erlangen-Nuremberg, Comprehensive Cancer Center Erlangen-EMN, Erlangen, Germany. [32] David Geffen School of Medicine, Department of Medicine Division of Hematology and Oncology, University of California at Los Angeles, Los Angeles, CA, USA. [33] Department of Biostatistics, Kansas University Medical Center, Kansas City, KS, USA. [34] Division of Genetics and Epidemiology, Institute of Cancer Research, London, UK. [35] Epidemiology Research Program, American Cancer Society, Atlanta, GA, USA. [36] Cancer Epidemiology & Intelligence Division, Cancer Council Victoria, Melbourne, Victoria, Australia. [37] Department of Epidemiology and Preventive Medicine, Monash University, Melbourne, Victoria, Australia. [38] Department of Health Science Research, Division of Epidemiology, Mayo Clinic, Rochester, MN, USA. [39] Department of Preventive Medicine, Keck School of Medicine, University of Southern California, Los Angeles, CA, USA. [40] Department of Oncology, Södersjukhuset, Stockholm, Sweden. [41] Department of Biostatistics & Epidemiology, University of Massachusetts, Amherst, Amherst, MA, USA. [42] Department of Clinical Genetics, St George's, University of London, London, UK. [43] Centre for Cancerbiomarkers, Department of Clinical Science, University of

Bergen, Bergen, Norway. [44] Department of Gynecology and Obstetrics, Haukeland University Hospital, Bergen, Norway. [45] Program in Genetic Epidemiology and Statistical Genetics, Harvard T.H. Chan School of Public Health, Boston, MA, USA. [46] Nuffield Department of Population Health, University of Oxford, Oxford, UK. [47] VIB Center for Cancer Biology, VIB, Leuven, Belgium. [48] Epidemiology Program, University of Hawaii Cancer Center, Honolulu, HI, USA. [49] Department of Epidemiology and Biostatistics, Memorial Sloan-Kettering Cancer Center, New York, NY, USA. [50] Department of Molecular Medicine and Surgery, Karolinska Institutet, Stockholm, Sweden. [51] Department of Clinical Genetics, Karolinska University Hospital, Stockholm, Sweden. [52] Department of Cancer Epidemiology and Prevention, M. Sklodowska-Curie Cancer Center, Oncology Institute, Warsaw, Poland. [53] Division of Epidemiology, Department of Medicine, Vanderbilt Epidemiology Center, Vanderbilt-Ingram Cancer Center, Vanderbilt University School of Medicine, Nashville, TN, USA. [54] Chronic Disease Epidemiology, Yale School of Public Health, New Haven, CT, USA. [55] Department of Anatomic Pathology, Moffitt Cancer Center & Research Institute, Tampa, FL, USA. [56] Institute of Cancer and Genomic Sciences, University of Birmingham, Birmingham, UK. [57] Precision Medicine, School of Clinical Sciences at Monash Health, Monash University, Clayton, Victoria, Australia. [58] Department of Women's and Children's Health, Karolinska Institutet, Stockholm, Sweden. [59] Department of Biochemistry and Molecular Medicine, University of California Davis, Davis, CA, USA. [60] School of Medicine and Public Health, University of Newcastle, Callaghan, New South Wales, Australia. [61] Harvard T.H. Chan School of Public Health, Boston, MA, USA. [62] Dana-Farber Cancer Institute, Boston, MA, USA. [63] Department of Gynaecology and Obstetrics, University Hospital Erlangen, Friedrich-Alexander University Erlangen-Nuremberg, Comprehensive Cancer Center Erlangen-EMN, Erlangen, Germany. [64] Center for Cancer Prevention (CPO-Peimonte), Turin, Italy. [65] Human Genetics Foundation (HuGeF), Turino, Italy. [66] Department of Obstetrics and Gynecology, School of Medicine and Public Health, University of Wisconsin, Madison, WI, USA. [67] Department of Population and Quantitative Health Sciences, Case Western Reserve University, Cleveland, OH, USA. [68] Division of Molecular Medicine, Pathology North, John Hunter Hospital, Newcastle, New South Wales, Australia. [69] Clinical Genetics, Karolinska Institutet, Stockholm, Sweden. [70] Division of Gynecologic Oncology, Department of Obstetrics and Gynaecology and Leuven Cancer Institute, University Hospitals Leuven, Leuven, Belgium. [71] Population Health Department, QIMR Berghofer Medical Research Institute, Brisbane, Queensland, Australia. [72] Department of Health Science Research, Division of Biomedical Statistics and Informatics, Mayo Clinic, Rochester, MN, USA. [73] State Key Laboratory of Oncogene and Related Genes & Department of Epidemiology, Shanghai Cancer Institute, Renji Hospital, Shanghai Jiaotong University School of Medicine, Shanghai, China.

**BIOS Consortium (Biobank-based Integrative Omics Study) – Author information**

**Management Team** Bastiaan T. Heijmans (chair)[1], Peter A.C. 't Hoen[2], Joyce van Meurs[3], Aaron Isaacs[4], Rick Jansen[5], Lude Franke[6].
**Cohort collection** Dorret I. Boomsma[7], René Pool[7], Jenny van Dongen[7], Jouke J. Hottenga[7] (Netherlands Twin Register); Marleen MJ van Greevenbroek[8], Coen D.A. Stehouwer[8], Carla J.H. van der Kallen[8], Casper G. Schalkwijk[8] (Cohort study on Diabetes and Atherosclerosis Maastricht); Cisca Wijmenga[6], Lude Franke[6], Sasha Zhernakova[6], Ettje F. Tigchelaar[6] (LifeLines Deep); P. Eline Slagboom[1], Marian Beekman[1], Joris Deelen[1], Diana van Heemst[9]

(Leiden Longevity Study); Jan H. Veldink[10], Leonard H. van den Berg[10] (Prospective ALS Study Netherlands); Cornelia M. van Duijn[4], Bert A. Hofman[11], Aaron Isaacs[4], André G. Uitterlinden[3] (Rotterdam Study).

**Data Generation** Joyce van Meurs (Chair)[3], P. Mila Jhamai[3], Michael Verbiest[3], H. Eka D. Suchiman[1], Marijn Verkerk[3], Ruud van der Breggen[1], Jeroen van Rooij[3], Nico Lakenberg[1]. Data management and computational infrastructure Hailiang Mei (Chair)[12], Maarten van Iterson[1], Michiel van Galen[2], Jan Bot[13], Dasha V. Zhernakova[6], Rick Jansen[5], Peter van 't Hof[12], Patrick Deelen[6], Irene Nooren[13], Peter A.C. 't Hoen[2], Bastiaan T. Heijmans[1], Matthijs Moed[1].

**Data Analysis Group** Lude Franke (Co-Chair)[6], Martijn Vermaat[2], Dasha V. Zhernakova[6], René Luijk[1], Marc Jan Bonder[6], Maarten van Iterson[1], Patrick Deelen[6], Freerk van Dijk[14], Michiel van Galen[2], Wibowo Arindrarto[12], Szymon M. Kielbasa[15], Morris A. Swertz[14], Erik. W van Zwet[15], Rick Jansen[5], Peter-Bram 't Hoen (Co-Chair)[2], Bastiaan T. Heijmans (Co-Chair)[1].

1. Molecular Epidemiology Section, Department of Medical Statistics and Bioinformatics, Leiden University Medical Center, Leiden, The Netherlands
2. Department of Human Genetics, Leiden University Medical Center, Leiden, The Netherlands
3. Department of Internal Medicine, ErasmusMC, Rotterdam, The Netherlands
4. Department of Genetic Epidemiology, ErasmusMC, Rotterdam, The Netherlands
5. Department of Psychiatry, VU University Medical Center, Neuroscience Campus Amsterdam, Amsterdam, The Netherlands
6. Department of Genetics, University of Groningen, University Medical Centre Groningen, Groningen, The Netherlands
7. Department of Biological Psychology, VU University Amsterdam, Neuroscience Campus Amsterdam, Amsterdam, The Netherlands
8. Department of Internal Medicine and School for Cardiovascular Diseases (CARIM), Maastricht University Medical Center, Maastricht, The Netherlands
9. Department of Gerontology and Geriatrics, Leiden University Medical Center, Leiden, The Netherlands
10. Department of Neurology, Brain Center Rudolf Magnus, University Medical Center Utrecht, Utrecht, The Netherlands
11. Department of Epidemiology, ErasmusMC, Rotterdam, The Netherlands
12. Sequence Analysis Support Core, Leiden University Medical Center, Leiden, The Netherlands
13. SURFsara, Amsterdam, the Netherlands
14. Genomics Coordination Center, University Medical Center Groningen, University of Groningen, Groningen, the Netherlands
15. Medical Statistics Section, Department of Medical Statistics and Bioinformatics, Leiden University Medical Center, Leiden, The Netherlands

eQTLGen Consortium – Author information

Mawussé Agbessi[1], Habibul Ahsan[2], Isabel Alves[1], Anand Andiappan[3], Wibowo Arindrarto[4], Philip Awadalla[1], Alexis Battle[5,6], Frank Beutner[7], Marc Jan Bonder[8,9,10], Dorret Boomsma[11], Mark Christiansen[12], Annique Claringbould[8], Patrick Deelen[8,13], Tõnu Esko[14], Marie-Julie Favé[1], Lude Franke[8], Timothy Frayling[15], Sina A. Gharib[16,12], Gregory Gibson[17], Bastiaan T. Heijmans[4], Gibran Hemani[18], Rick Jansen[19], Mika Kähönen[20,21], Anette Kalnapenkis[14], Silva Kasela[14], Johannes Kettunen[22], Yungil Kim[6,23], Holger Kirsten[24], Peter Kovacs[25], Knut Krohn[26], Jaanika Kronberg-Guzman[14], Viktorija Kukushkina[14], Zoltan Kutalik[27], Bernett Lee[3], Terho Lehtimäki[28,29], Markus Loeffler[24], Urko M. Marigorta[17], Hailang Mei[4], Lili Milani[14], Grant W. Montgomery[30], Martina Müller-Nurasyid[31,32,33], Matthias Nauck[34], Michel Nivard[11], Brenda Penninx[19], Markus Perola[35], Natalia Pervjakova[14], Brandon L. Pierce[2], Joseph Powell[36], Holger Prokisch[37,38], Bruce M. Psaty[12,39,40], Olli T. Raitakari[41,42], Samuli Ripatti[43], Olaf Rotzschke[3], Sina Rüeger[27], Ashis Saha[6], Markus Scholz[24], Katharina Schramm[31,32], Ilkka Seppälä[28,29], Eline P. Slagboom[4], Coen D.A. Stehouwer[44], Michael Stumvoll[45], Patrick Sullivan[46], Peter-Bram ‘t Hoen[47], Alexander Teumer[48], Joachim Thiery[49], Lin Tong[2], Anke Tönjes[45], Jenny van Dongen[11], Maarten van Iterson[4], Joyce van Meurs[50], Jan H. Veldink[51], Joost Verlouw[50], Peter M. Visscher[30], Uwe Völker[52], Urmo Võsa[8,14], Harm-Jan Westra[8], Cisca Wijmenga[8], Hanieh Yaghootkar[15], Jian Yang[30,53], Biao Zeng[17], Futao Zhang[30]

Author list is ordered alphabetically.

1. Computational Biology, Ontario Institute for Cancer Research, Toronto, Canada
2. Department of Public Health Sciences, University of Chicago, Chicago, United States of America
3. Singapore Immunology Network, Agency for Science, Technology and Research, Singapore, Singapore
4. Department of Biomedical Data Sciences, Leiden University Medical Center, Leiden, The Netherlands
5. Department of Biomedical Engineering, Johns Hopkins University, Baltimore, United States of America
6. Department of Computer Science, Johns Hopkins University, Baltimore, United States of America
7. Heart Center Leipzig, Universität Leipzig, Leipzig, Germany
8. Department of Genetics, University Medical Centre Groningen, Groningen, The Netherlands
9. European Molecular Biology Laboratory, European Bioinformatics Institute, Wellcome Genome Campus, Hinxton, United Kingdom
10. Genome Biology Unit, European Molecular Biology Laboratory, Heidelberg, Germany
11. Department of Biological Psychology, Vrije Universiteit Amsterdam, Amsterdam, The Netherlands
12. Cardiovascular Health Research Unit, University of Washington, Seattle, United States of America
13. Genomics Coordination Center, University Medical Centre Groningen, Groningen, The Netherlands

14. Estonian Genome Center, Institute of Genomics, University of Tartu, Tartu 51010, Estonia
15. Exeter Medical School, University of Exeter, Exeter, United Kingdom
16. Department of Medicine, University of Washington, Seattle, United States of America
17. School of Biological Sciences, Georgia Tech, Atlanta, United States of America
18. MRC Integrative Epidemiology Unit, University of Bristol, Bristol, United Kingdom
19. Department of Psychiatry and Amsterdam Neuroscience, Amsterdam UMC, Vrije Universiteit, Amsterdam, the Netherlands
20. Department of Clinical Physiology, Tampere University, Tampere, Finland
21. Department of Clinical Physiology, Finnish Cardiovascular Research Center – Tampere, Faculty of Medicine and Life Sciences, University of Tampere, Tampere, Finland
22. Centre for Life Course Health Research, University of Oulu, Oulu, Finland
23. Genetics and Genomic Science Department, Icahn School of Medicine at Mount Sinai, New York, United States of America
24. Institut für Medizinische Informatik, Statistik und Epidemiologie, LIFE – Leipzig Research Center for Civilization Diseases, Universität Leipzig, Leipzig, Germany
25. IFB Adiposity Diseases, Universität Leipzig, Leipzig, Germany
26. Interdisciplinary Center for Clinical Research, Faculty of Medicine, Universität Leipzig, Leipzig, Germany
27. Institute of Social and Preventive Medicine, Lausanne University Hospital, Lausanne, Switzerland
28. Department of Clinical Chemistry, Fimlab Laboratories, Tampere, Finland
29. Department of Clinical Chemistry, Finnish Cardiovascular Research Center - Tampere, Faculty of Medicine and Life Sciences, University of Tampere, Tampere, Finland
30. Institute for Molecular Bioscience, University of Queensland, Brisbane, Australia
31. Institute of Genetic Epidemiology, Helmholtz Zentrum München - German Research Center for Environmental Health, Neuherberg, Germany
32. Department of Medicine I, University Hospital Munich, Ludwig Maximilian's University, München, Germany
33. DZHK (German Centre for Cardiovascular Research), partner site Munich Heart Alliance, Munich, Germany
34. Institute of Clinical Chemistry and Laboratory Medicine, University Medicine Greifswald, Greifswald, Germany
35. National Institute for Health and Welfare, University of Helsinki, Helsinki, Finland
36. Garvan Institute of Medical Research, Garvan-Weizmann Centre for Cellular Genomics, Sydney, Australia
37. Institute of Human Genetics, Helmholtz Zentrum München, Neuherberg, Germany
38. Institute of Human Genetics, Technical University Munich, Munich, Germany
39. Departments of Epidemiology, Medicine, and Health Services, University of Washington, Seattle, United States of America
40. Kaiser Permanente Washington Health Research Institute, Seattle, WA, United States of America
41. Department of Clinical Physiology and Nuclear Medicine, Turku University Hospital, Turku, Finland
42. Research Centre of Applied and Preventive Cardiovascular Medicine, University of Turku, Turku, Finland
43. Statistical and Translational Genetics, University of Helsinki, Helsinki, Finland

44. Department of Internal Medicine, Maastricht University Medical Centre, Maastricht, The Netherlands
45. Department of Medicine, Universität Leipzig, Leipzig, Germany
46. Department of Medical Epidemiology and Biostatistics, Karolinska Institutet, Stockholm, Sweden
47. Center for Molecular and Biomolecular Informatics, Radboud Institute for Molecular Life Sciences, Radboud University Medical Center Nijmegen, Nijmegen, The Netherlands
48. Institute for Community Medicine, University Medicine Greifswald, Greifswald, Germany
49. Institute for Laboratory Medicine, LIFE – Leipzig Research Center for Civilization Diseases, Universität Leipzig, Leipzig, Germany
50. Department of Internal Medicine, Erasmus Medical Centre, Rotterdam, The Netherlands
51. Department of Neurology, University Medical Center Utrecht, Utrecht, The Netherlands
52. Interfaculty Institute for Genetics and Functional Genomics, University Medicine Greifswald, Greifswald, Germany
53. Institute for Advanced Research, Wenzhou Medical University, Wenzhou, Zhejiang 325027, China

**Acknowledgements**

We thank the 23andMe research participants who contributed to this study

K. Ström is acknowledged for collection of samples for the single cell project. This work was supported by grants from the European Research Council ERC Starting Grant, the Swedish Research Council, and Kjell och Märta Beijers Stiftelse to L.A.F and by the Hjärnfonden, Swedish Cancer Society, the Swedish Research Council, Alzheimerfonden and the Foundation for Polish Science under the International Research Agendas Programme to J.P.D. Sequencing was performed by the SNP&SEQ Technology Platform in Uppsala. The facility is part of the National Genomics Infrastructure (NGI) Sweden and Science for Life Laboratory. The SNP&SEQ Platform is also supported by the Swedish Research Council and the Knut and Alice Wallenberg Foundation.

Stephen J Chanock is supported by the NIH Intramural Research Program. Rong Li is supported by the National Institute of Health (R35-GM118172)

Po-Ru Loh is supported by NIH grant DP2-ES030554, a Burroughs Wellcome Fund Career Award at the Scientific Interfaces, a Glenn Foundation for Medical Research and AFAR Grants for Junior Faculty award, and the Next Generation Fund at the Broad Institute of MIT and Harvard.

J.C.U. is supported by a National Institutes of Health training grant (5T32 GM007226-43). J.C.U acknowledges analytical help and advice from Christian Benner, Liam Abbott, Masahiro Kanai, Erik Bao, Caleb Lareau

**Lung Cancer Consortium**

Transdisciplinary Research for Cancer in Lung (TRICL) of the International Lung Cancer Consortium (ILCCO) was supported by grants U19-CA148127 and CA148127S1. ILCCO data harmonization is supported by the Cancer Care Ontario Research Chair of Population Studies to R.J.H. and the Lunenfeld-Tanenbaum Research Institute, Sinai Health System. The TRICL-ILCCO OncoArray was supported by in-kind genotyping by the Centre for Inherited Disease Research (26820120008i-0-26800068-1). IARC acknowledges and thanks V. Gaborieau, M. Foll, L. Fernandez-Cuesta, P. Chopard, T. Delhomme and A. Chabrier for their technical assistance in this project. The authors would like to thank the staff at the Respiratory Health Network Tissue Bank of the FRQS for their valuable assistance with the lung eQTL data set at Laval University. The lung eQTL study at Laval University was supported by the Fondation de l'Institut Universitaire de Cardiologie et de Pneumologie de Québec, the Respiratory Health Network of the FRQS and the Canadian Institutes of Health Research (MOP-123369).

The CAPUA study was supported by FIS-FEDER/Spain grant numbers FIS-01/310, FIS-PI03-0365, and FIS-07-BI060604, FICYT/Asturias grant numbers FICYT PB02-67 and FICYT IB09-133, and the University Institute of Oncology (IUOPA), of the University of Oviedo and the Ciber de Epidemiologia y Salud Pública. CIBERESP, SPAIN.

The work performed in the CARET study was supported by the National Institute of Health / National Cancer Institute: UM1 CA167462 (PI: Goodman), National Institute of Health UO1-

CA6367307 (PIs Omen, Goodman); National Institute of Health R01 CA111703 (PI Chen), National Institute of Health 5R01CA151989-01A1(PI Doherty).

The Liverpool Lung project is supported by the Roy Castle Lung Cancer Foundation.

The Harvard Lung Cancer Study was supported by the NIH (National Cancer Institute) grants CA092824, CA090578, CA074386.

The Multiethnic Cohort Study was partially supported by NIH Grants CA164973, CA033619, CA63464 and CA148127.

The work performed in MSH-PMH study was supported by The Canadian Cancer Society Research Institute (020214), Ontario Institute of Cancer and Cancer Care Ontario Chair Award to R.J.H. and G.L. and the Alan Brown Chair and Lusi Wong Programs at the Princess Margaret Hospital Foundation.

NJLCS was funded by the State Key Program of National Natural Science of China (81230067), the National Key Basic Research Program Grant (2011CB503805), the Major Program of the National Natural Science Foundation of China (81390543).

The Norway study was supported by Norwegian Cancer Society, Norwegian Research Council.

The Shanghai Cohort Study (SCS) was supported by National Institutes of Health R01 CA144034 (PI: Yuan) and UM1 CA182876 (PI: Yuan).

The Singapore Chinese Health Study (SCHS) was supported by National Institutes of Health R01 CA144034 (PI: Yuan) and UM1 CA182876 (PI: Yuan).

The work in TLC study has been supported in part the James & Esther King Biomedical Research Program (09KN-15),National Institutes of Health Specialized Programs of Research Excellence (SPORE) Grant (P50 CA119997), and by a Cancer Center Support Grant (CCSG) at the H. Lee Moffitt Cancer Center and Research Institute, an NCI designated Comprehensive Cancer Center (grant number P30-CA76292).

The Vanderbilt Lung Cancer Study – BioVU dataset used for the analyses described was obtained from Vanderbilt University Medical Center's BioVU, which is supported by institutional funding, the 1S10RR025141-01 instrumentation award, and by the Vanderbilt CTSA grant UL1TR000445 from NCATS/NIH. Dr. Aldrich was supported by NIH/National Cancer Institute K07CA172294 (PI:Aldrich).

The Copenhagen General Population Study (CGPS) was supported by the Chief Physician Johan Boserup and Lise Boserup Fund, the Danish Medical Research Council and Herlev Hospital.

The NELCS study: Grant Number P20RR018787 from the National Center for Research Resources (NCRR), a component of the National Institutes of Health (NIH).

The Kentucky Lung Cancer Research Initiative was supported by the Department of Defense [Congressionally Directed Medical Research Program, U.S. Army Medical Research and Materiel Command Program] under award number: 10153006 (W81XWH-11-1-0781). Views and opinions of, and endorsements by the author(s) do not reflect those of the US Army or the Department of Defense. This research was also supported by unrestricted infrastructure funds from the UK Center for Clinical and Translational Science, NIH grant UL1TR000117 and Markey Cancer Center NCI Cancer Center Support Grant (P30 CA177558) Shared Resource Facilities: Cancer Research Informatics, Biospecimen and Tissue Procurement, and Biostatistics and Bioinformatics.

The M.D. Anderson Cancer Center study was supported in part by grants from the NIH (P50 CA070907, R01 CA176568) (to X. Wu), Cancer Prevention & Research Institute of Texas (RP130502) (to X. Wu), and The University of Texas MD Anderson Cancer Center institutional support for the Center for Translational and Public Health Genomics.

The study in Lodz center was partially funded by Nofer Institute of Occupational Medicine, under task NIOM 10.13: Predictors of mortality from non-small cell lung cancer - field study.

Genetic sharing analysis was funded by NIH grant CA194393

The work to assemble the FTND GWAS meta-analysis was supported by the National Institutes of Health (NIH), National Institute on Drug Abuse (NIDA) grant number R01 DA035825 (Principal Investigator [PI]: DBH). The study populations included COGEND (dbGaP phs000092.v1.p1 and phs000404.v1.p1), COPDGene (dbGaP phs000179.v3.p2), deCODE Genetics, EAGLE (dbGaP phs000093.vs.p2), and SAGE. dbGaP phs000092.v1.p1).

**The PRACTICAL Consortium**

Genotyping of the OncoArray was funded by the US National Institutes of Health (NIH) [U19 CA 148537 for ELucidating Loci Involved Prostate cancer SuscEptibility (ELLIPSE) project and X01HG007492 to the Center for Inherited Disease Research (CIDR) under contract number HHSN268201200008I]. Additional analytic support was provided by NIH NCI U01 CA188392 (PI: Schumacher). The PRACTICAL consortium was supported by Cancer Research UK Grants C5047/A7357, C1287/A10118, C1287/A16563, C5047/A3354, C5047/A10692, C16913/A6135, European Commission's Seventh Framework Programme grant agreement n° 223175 (HEALTH-F2-2009-223175), and The National Institute of Health (NIH) Cancer Post-Cancer GWAS initiative grant: No. 1 U19 CA 148537-01 (the GAME-ON initiative). We would also like to thank the following for funding support: The Institute of Cancer Research and The Everyman Campaign, The Prostate Cancer Research Foundation, Prostate Research Campaign UK (now Prostate Action), The Orchid Cancer Appeal, The National Cancer Research Network UK, The National Cancer Research Institute (NCRI) UK. We are grateful for support of NIHR funding to the NIHR Biomedical Research Centre at The Institute of Cancer Research and The Royal Marsden NHS Foundation Trust.

**The Breast Cancer Association Consortium**

The breast cancer genome-wide association analyses were supported by the Government of Canada through Genome Canada and the Canadian Institutes of Health Research, the 'Ministère de l'Économie, de la Science et de l'Innovation du Québec' through Genome Québec and grant PSR-SIIRI-701, The National Institutes of Health (U19 CA148065, X01HG007492), Cancer Research UK (C1287/A10118, C1287/A16563, C1287/A10710) and The European Union (HEALTH-F2-2009-223175 and H2020 633784 and 634935). All studies and funders are listed Michailidou et al (2017) [PMID 29059683].

**Consortium of Investigators of Modifiers of BRCA1/2 (CIMBA)**

The CIMBA data management and data analysis were supported by Cancer Research – UK grants C12292/A20861, C12292/A11174. ACA is a Cancer Research -UK Senior Cancer Research Fellow. GCT and ABS are NHMRC Research Fellows. iCOGS: the European Community's Seventh Framework Programme under grant agreement n° 223175 (HEALTH-F2-2009-223175) (COGS), Cancer Research UK (C1287/A10118, C1287/A 10710, C12292/A11174, C1281/A12014, C5047/A8384, C5047/A15007, C5047/A10692, C8197/A16565), the National Institutes of Health (CA128978) and Post-Cancer GWAS initiative (1U19 CA148537, 1U19 CA148065 and 1U19 CA148112 - the GAME-ON initiative), the Department of Defence (W81XWH-10-1-0341), the Canadian Institutes of Health Research (CIHR) for the CIHR Team Familial Risks of Breast Cancer (CRN-87521), and the Ministry of Economic Development, Innovation and Export Trade (PSR-SIIRI-701), Komen Foundation for the Cure, the Breast Cancer Research Foundation, and the Ovarian Cancer Research Fund. The PERSPECTIVE project was supported by the Government of Canada through Genome Canada and the Canadian Institutes of Health Research, the Ministry of Economy, Science and Innovation through Genome Québec, and The Quebec Breast Cancer Foundation.

**Colorectal Cancer UK GWAS**

At the Institute of Cancer Research, this work was supported by Cancer Research UK (C1298/A25514). Additional support was provided by the National Cancer Research Network. The COIN and COIN-B trials were funded by Cancer Research UK and the Medical Research Council and were conducted with the support of the National Institute of Health Research Cancer Research Network. COIN and COIN-B translational studies were supported by the Bobby Moore Fund from Cancer Research UK, Tenovus, the Kidani Trust, Cancer Research Wales and the National Institute for Social Care and Health Research Cancer Genetics Biomedical Research Unit (2011–2014). For control sample genotypes, we acknowledge the PRACTICAL consortium support from Cancer Research UK Grants

C5047/A7357, C1287/A10118, C1287/A16563, C5047/A3354, C5047/A10692, C16913/A6135, European Commission's Seventh Framework Programme grant agreement n° 223175 (HEALTH-F2-2009-223175), and The National Institute of Health (NIH) Cancer Post-Cancer GWAS initiative grant: No. 1 U19 CA 148537-01 (the GAME-ON initiative).

In Birmingham, funding was provided by Cancer Research UK (C6199/A16459). In Oxford additional funding was provided by the Oxford Comprehensive Biomedical Research Centre and the EU FP7 CHIBCHA grant. Core infrastructure support to the Wellcome Trust Centre for Human Genetics, Oxford was provided by grant (090532/Z/09/Z). We are grateful to many colleagues within UK Clinical Genetics Departments (for CORGI) and to many collaborators who participated in the VICTOR, QUASAR2 and SCOT trials. We acknowledge use of genotype data from the British 1958 Birth Cohort DNA collection, which was funded by the Medical Research Council Grant G0000934 and the Wellcome Trust Grant 068545/Z/02. A full list of the investigators who contributed to the generation of the data is available from http://www.wtccc.org.uk.

In Edinburgh, the work was supported by Programme Grant funding from Cancer Research UK (C348/A18927) and by MRC Centre Grant funding (U127527202 and U127527198 from 1/4/18) to the MRC Human Genetics Unit. Infrastructure and staffing funding is acknowledged from the Edinburgh CRUK Cancer Research Centre grant. The Lothian Birth Cohort studies are funded by Age UK (Disconnected Mind project) and the Biotechnology and Biological Sciences Research Council (grant no. BB/F019394/1). Genotyping of the GS:SFHS samples was carried out by the Edinburgh Clinical Research Facility, University of Edinburgh, and was funded by the Medical Research Council UK and the Wellcome Trust (Wellcome Trust Strategic Award 'STratifying Resilience and Depression Longitudinally' (STRADL), Reference 104036/Z/14/Z). GS:SFHS received core support from the Scottish Executive Health Department, Chief Scientist Office, grant number CZD/16/6. The MRC provides core funding to the QTL in Health and Disease research program at the MRC HGU, IGMM, University of Edinburgh. The inclusion of UK Biobank cases and controls was conducted under the UK Biobank Resource Application Number 7441.

**The Endometrial Cancer Association Consortium**

The authors thank the many individuals who participated in this study and the numerous institutions and their staff who supported recruitment.

The iCOGS and OncoArray endometrial cancer analysis were supported by NHMRC project grants [ID#1031333 & ID#1109286] Funding for the iCOGS infrastructure came from: the European Community's Seventh Framework Programme under grant agreement no 223175 [HEALTH-F2-2009-223175] [COGS], Cancer Research UK [C1287/A10118, C1287/A 10710, C12292/A11174, C1281/A12014, C5047/A8384, C5047/A15007, C5047/A10692, C8197/A16565], the National Institutes of Health [CA128978] and Post-Cancer GWAS initiative [1U19 CA148537, 1U19 CA148065 and 1U19 CA148112 - the GAME-ON initiative], the Department of Defence [W81XWH-10-1-0341], the Canadian Institutes of Health Research [CIHR] for the CIHR Team in Familial Risks of Breast Cancer, Komen Foundation for the Cure, the Breast Cancer Research Foundation, and the Ovarian Cancer Research Fund. OncoArray genotyping of ECAC cases was performed with the generous assistance of the Ovarian Cancer Association Consortium (OCAC). We particularly thank the efforts of Cathy Phelan. The OCAC OncoArray genotyping project was funded through grants from the US National Institutes of Health (CA1X01HG007491-01, U19-CA148112, R01-CA149429 and R01-CA058598); Canadian Institutes of Health Research (MOP-86727); and the Ovarian Cancer Research Fund. CIDR genotyping for the Oncoarray was conducted under contract 268201200008I. OncoArray genotyping of the BCAC controls was funded by Genome Canada Grant GPH-129344, NIH Grant U19 CA148065, and Cancer UK Grant C1287/A16563.

Stage 1 and stage 2 case genotyping was supported by the NHMRC [ID#552402, ID#1031333]. Control data were generated by the Wellcome Trust Case Control Consortium (WTCCC), and a full list of the investigators who contributed to the generation of the data is available from the WTCCC website. We acknowledge use of DNA from the British 1958 Birth Cohort collection, funded by the Medical Research Council grant G0000934 and the Wellcome Trust grant 068545/Z/02 - funding for this project was provided by the Wellcome Trust under award 085475. NSECG was supported by the EU FP7 CHIBCHA grant, Wellcome Trust Centre for Human Genetics Core Grant 090532/Z/09Z, and CORGI was funded by Cancer Research UK. We thank Nick Martin, Dale Nyholt and Anjali Henders for access to GWAS data from QIMR Controls. Recruitment of the QIMR controls was supported by the NHMRC. The University of Newcastle, the Gladys M Brawn Senior Research Fellowship scheme, The Vincent Fairfax Family Foundation, the Hunter Medical Research Institute and the Hunter Area Pathology Service all contributed towards the costs of establishing the Hunter Community Study. The WHI program is funded by the National Heart, Lung, and Blood Institute, the US National Institutes of Health and the US Department of Health and Human Services (HHSN268201100046C, HHSN268201100001C, HHSN268201100002C, HHSN268201100003C, HHSN268201100004C and HHSN271201100004C). This work was also funded by NCI U19 CA148065-01. This research has been conducted using the UK Biobank Resource under applications 5122 and 9797.

ANECS recruitment was supported by project grants from the NHMRC [ID#339435], The Cancer Council Queensland [ID#4196615] and Cancer Council Tasmania [ID#403031 and ID#457636]. SEARCH recruitment was funded by a programme grant from Cancer Research UK [C490/A10124]. The Bavarian Endometrial Cancer Study (BECS) was partly funded by the ELAN fund of the University of Erlangen. The Hannover-Jena Endometrial Cancer Study was partly supported by the Rudolf Bartling Foundation. The Leuven Endometrium Study (LES) was supported by the Verelst Foundation for endometrial cancer. The Mayo Endometrial Cancer Study (MECS) and Mayo controls (MAY) were supported by grants from the National Cancer Institute of United States Public Health Service [R01 CA122443, P30 CA15083, P50 CA136393, and GAME-ON the NCI Cancer Post-GWAS Initiative U19 CA148112], the Fred C and Katherine B Andersen Foundation, the Mayo Foundation, and the Ovarian Cancer Research Fund with support of the Smith family, in memory of Kathryn Sladek Smith. MoMaTEC received financial support from a Helse Vest Grant, the University of Bergen, Melzer Foundation, The Norwegian Cancer Society (Harald Andersens legat), The Research Council of Norway and Haukeland University Hospital. The Newcastle Endometrial Cancer Study (NECS) acknowledges contributions from the University of Newcastle, The NBN Children's Cancer Research Group, Ms Jennie Thomas and the Hunter Medical Research Institute. RENDOCAS was supported through the regional agreement on medical training and clinical research (ALF) between Stockholm County Council and Karolinska Institutet [numbers: 20110222, 20110483, 20110141 and DF 07015], The Swedish Labor Market Insurance [number 100069] and The Swedish Cancer Society [number 11 0439]. The Cancer Hormone Replacement Epidemiology in Sweden Study (CAHRES, formerly called The Singapore and Swedish Breast/Endometrial Cancer Study; SASBAC) was supported by funding from the Agency for Science, Technology and Research of Singapore (A*STAR), the US National Institutes of Health and the Susan G. Komen Breast Cancer Foundation.

The Nurses' Health Study (NHS) is supported by the NCI, NIH Grants Number UM1 CA186107, P01 CA087969, R01 CA49449, 1R01 CA134958, and 2R01 CA082838. The authors thank the participants and staff of the Nurses' Health Study for their valuable contributions as well as the following state cancer registries for their help: AL, AZ, AR, CA, CO, CT, DE, FL, GA, ID, IL, IN, IA, KY, LA, ME, MD, MA, MI, NE, NH, NJ, NY, NC, ND, OH, OK, OR, PA, RI, SC, TN, TX, VA, WA, WY. The authors assume full responsibility for analyses and interpretation of these data. The authors also thank Channing Division of Network Medicine, Department of Medicine, Brigham and Women's Hospital and Harvard Medical School. Finally, the authors also acknowledge Pati Soule and Hardeep Ranu for their laboratory assistance. The Connecticut Endometrial Cancer Study was supported by NCI, NIH Grant Number RO1CA98346. The Fred Hutchinson Cancer Research Center (FHCRC) is supported by NCI, NIH Grant Number NIH RO1 CA105212, RO1 CA 87538, RO1 CA75977, RO3 CA80636, NO1 HD23166, R35 CA39779, KO5 CA92002 and funds from the Fred Hutchinson Cancer Research Center. The Multiethnic Cohort Study (MEC) is supported by the NCI, NHI Grants Number CA54281, CA128008 and 2R01 CA082838. The California Teachers Study (CTS) is supported by NCI, NIH Grant Number 2R01 CA082838, R01 CA91019 and R01 CA77398, and contract 97-10500 from the California Breast Cancer Research Fund. The Polish Endometrial Cancer Study (PECS) is supported by the Intramural Research Program of the NCI. The Prostate, Lung, Colorectal, and Ovarian Cancer Screening Trial (PLCO) is supported by the Extramural and the Intramural Research Programs of the NCI.

**Testicular Germ Cell Tumour GWAS**

We thank the subjects with TGCT and the clinicians involved in their care for participation in this work. We thank the patients and all clinicians forming part of the UK Testicular Cancer Collaboration (UKTCC) for their participation in this work. This work makes use of data generated by the Wellcome Trust Case Control Consortium 2 (WTCCC2). We acknowledge the contribution of E. Rapley, M. Stratton and K Litchfield to the generation of the GWAS data. This study was supported by the Movember Foundation and the Institute of Cancer Research. Additional support was provided as follows: genotyping of the OncoArray was funded by the US National Institutes of Health (NIH) [U19 CA 148537 for ELucidating Loci Involved in Prostate cancer SuscEptibility (ELLIPSE) project and X01HG007492 to the Center for Inherited Disease Research (CIDR) under contract number HHSN268201200008I]; additional analytic support was provided by NIH NCI U01 CA188392 (PI: Schumacher); the PRACTICAL consortium was supported by Cancer Research UK Grants C5047/A7357, C1287/A10118, C1287/A16563, C5047/A3354, C5047/A10692, C16913/A6135, European Commission's Seventh Framework Programme grant agreement n° 223175 (HEALTH-F2-2009-223175), and The National Institute of Health (NIH) Cancer Post-Cancer GWAS initiative grant: No. 1 U19 CA 148537-01 (the GAME-ON initiative).
